# Innovations in Alginate Catabolism Leading to Heterotrophy and Adaptive Evolution of Diatoms

**DOI:** 10.1101/2024.08.28.610029

**Authors:** Zeng Hao Lim, Peng Zheng, Christopher Quek, Minou Nowrousian, Finn L. Aachmann, Gregory Jedd

## Abstract

A major goal of evolutionary biology is to identify the genetic basis for the emergence of adaptive traits. Diatoms are ancestrally photosynthetic microalgae. However, in the genus *Nitzschia*, loss of photosynthesis led to a group of free-living secondary heterotrophs whose manner of energy acquisition is unclear. Here, we sequence the genome of the non-photosynthetic diatom *Nitzschia* sing1 and identify the genetic basis for its catabolism of the brown seaweed cell wall polysaccharide alginate. *N*. sing1 obtained an endolytic alginate lyase enzyme by horizontal gene transfer (HGT) from a marine bacterium. Subsequent gene duplication and transposition led to 91 genes in three distinct gene families. One family retains the ancestral endolytic enzyme function. By contrast, the two others underwent domain duplication, gain, loss, rearrangement, and mutation to encode novel functions that can account for oligosaccharide import through the endomembrane system and the exolytic production of alginate monosaccharides. Together, our results show how a single HGT event followed by substantial gene duplication and neofunctionalization led to alginate catabolism and access to a new ecological niche.

**Highlights:**

- *N*. sing1 acquired an alginate lyase (ALY) gene by horizontal gene transfer from a marine bacterium
- This founding gene expanded and diversified to comprise 3 major families across 30 loci
- Derived functions account for alginate import and processing into monomers
- Domain duplication, gain, loss, mutation, and *de novo* sequence evolution underlie ALY gene neofunctionalization

Graphical Abstract

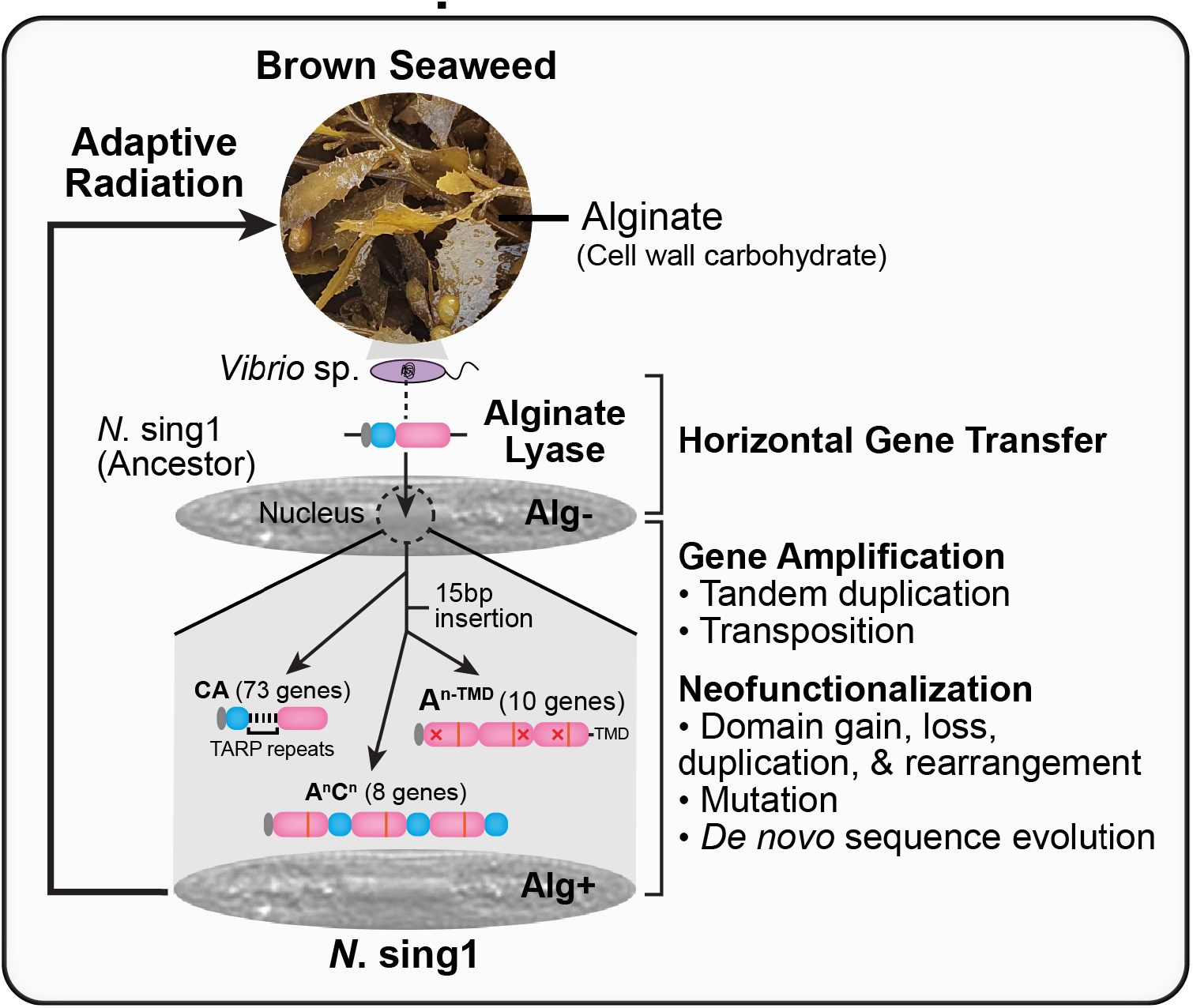

## Introduction

How organisms obtain energy is a fundamental determinant of their form, function, and evolutionary trajectory. Photoautotrophs obtain energy from sunlight, while heterotrophs depend on chemical energy derived from these primary producers. Transitions between trophic strategies constitute major evolutionary steps that can lead to adaptive radiation. A prime example is the acquisition of photosynthesis through endosymbiosis between a eukaryotic heterotroph and a photosynthetic cyanobacterium that occurred around 1.5 billion years ago^1^. This key event led to the emergence of land plants, green and red algae, and glaucophytes (Archaeplastida). Subsequent endosymbiosis between a green or red alga and eukaryotic heterotroph led to distinct algal lineages, which together with members of the Archaeplastida comprise the known eukaryotic photoautotrophs (reviewed in^2–5^).

While multiple endosymbiotic transitions and the diversity of eukaryotic photoautotrophs attest to the advantages of this trophic strategy, loss of photosynthesis leading to secondary heterotrophs has also occurred in each major photosynthetic lineage. In many cases, this involves a transition to parasitism and narrowing of the habitat range. Such transitions have occurred in flowering plants^6,7^, red^8^ and green^9^ algae, and apicomplexans such as *Plasmodium* and *Toxoplasma*^10,11^. Loss of photosynthesis has also led to many free-living secondary heterotrophs. This has occurred in the green^12–15^ and red^16^ algae, cryptophytes^17,18^, euglenids^19^, dinoflagellates^20^, colpodellids^21^, chrysophytes^22,23^ and diatoms^24,25^. Here, mechanisms for nutrient uptake include phagotrophy and osmotrophy. However, in most cases, the genetic and physiological basis for adaptation to obligate heterotrophy is poorly understood.

In the microalgal diatoms^26–28^, loss of photosynthesis in the genus *Nitzschia* led to free-living heterotrophs that occupy the nutrient-rich waters of the intertidal zone. These colorless or apochlorotic diatoms have been isolated from decaying plant material and the surface of green, red and brown algae^24,25,29–32^. Moreover, they have been shown to grow on sole carbon sources consisting of cellulose^33^, the red algal cell wall polysaccharides carrageenan and agarose^29,31–33^, and the brown seaweed cell wall polysaccharide alginate^32^. Genome sequences have revealed genetic signatures of apochlorotic diatoms. A β-ketoadipate pathway for metabolism of lignin-derived aromatic compounds and rewiring of mitochondrial glycolysis have been implicated in *N.* Nitz4^34^, while expansion and diversification of solute transporters, carbohydrate-active enzymes, and a unique secretome have been documented in *N. putrida*^35^.

Here, we sequence the genome of the apochlorotic diatom *N*. sing1 and show that its ancestor acquired a Polysaccharide Lyase 7 (PL7) family alginate lyase (ALY) gene by horizontal gene transfer (HGT) from a marine bacterium. This founder gene went on to expand through a combination of tandem gene duplication and transposition. Subsequent diversification of the paralogs gave rise to three major families comprising 91 genes. One *N.* sing1 ALY family retains the original endolytic function. By contrast, the two others underwent domain duplication, gain, loss, mutation, and substantial rearrangement to encode new functions that can account for alginate oligosaccharide import through the endomembrane system and their conversion into monomers. Thus, a full alginate catabolic pathway appears to have originated through neofunctionalization of paralogs derived from a single gene obtained by HGT. ALY genes are absent from the apochlorotic diatoms *N*. *putrida* and *N*. Nitz4, suggesting a high degree of ecophysiological diversity within the apochlorotic lineage. Together, our data show how HGT, coupled with gene duplication and neofunctionalization, led to the evolution of a complex metabolic capability and adaptive radiation onto brown seaweed habitats.

## Results

### N. sing1 obtained an alginate lyase enzyme by horizontal gene transfer

To investigate the genetic basis for obligate heterotrophy in diatoms, we sequenced, assembled, and annotated the genome of *Nitzschia* sing1 (*N*. sing1) (see Materials and Methods). The assembly spans 40.35 Mbp and is predicted to encode 15,542 protein-coding genes (Supplementary Table 1). Analysis of genome heterozygosity indicates that vegetative *N*. sing1 cells are diploid (Supplementary Fig. 1), as are other diatoms^35–40^. Putative telomeric repeats were found at both ends of one contig and at one end of another 22 contigs (Supplementary Table 2), making it likely that the *N*. sing1 genome consists of at least 12 chromosomes. BUSCO (Benchmarking Universal Single-Copy Orthologs) analysis shows that the assembly is relatively complete and comparable to that of other sequenced diatoms (Supplementary Table 1).

Because apochlorotic diatoms grow on diverse seaweed-derived polysaccharides, we searched the *N*. sing1 genome for carbohydrate-active enzymes (CAZymes)^41^. Here, we compared the CAZyme profile of *N*. sing1 to the two sequenced apochlorotic species (*N*. *putrida*^35^ *and N*. Nitz4^34^), five photosynthetic diatoms^36–40^, and five other species in the SAR (stramenopiles, alveolates, and rhizaria) supergroup^42–47^. These data show that *N*. sing1 encodes an unusually large number (91) of genes containing at least one Polysaccharide Lyase 7 (PL7) family alginate lyase domain (Fig. 1a, b, Supplementary Fig. 2 and Supplementary Table 3). PL7 family alginate lyases are well-characterized enzymes that cleave the glycosidic bonds between alginate sugar residues through a β-elimination mechanism^48–50^. Besides the PL7 catalytic domain, a large majority of *N*. sing1 alginate lyase genes (ALYs) also encode Carbohydrate-Binding Module 32 (CBM32) domains (Fig. 1b and Supplementary Table 4), which have been shown to bind various carbohydrate moieties^51,52^ and are implicated as modifiers of alginate lyase specificity^53–55^. Interestingly, neither the photosynthetic nor other apochlorotic diatoms encode any ALY genes (Fig. 1a and Supplementary Fig. 2), suggesting that the acquisition and expansion of this gene family is unique to the *N*. sing1 lineage.

**Figure 1.**
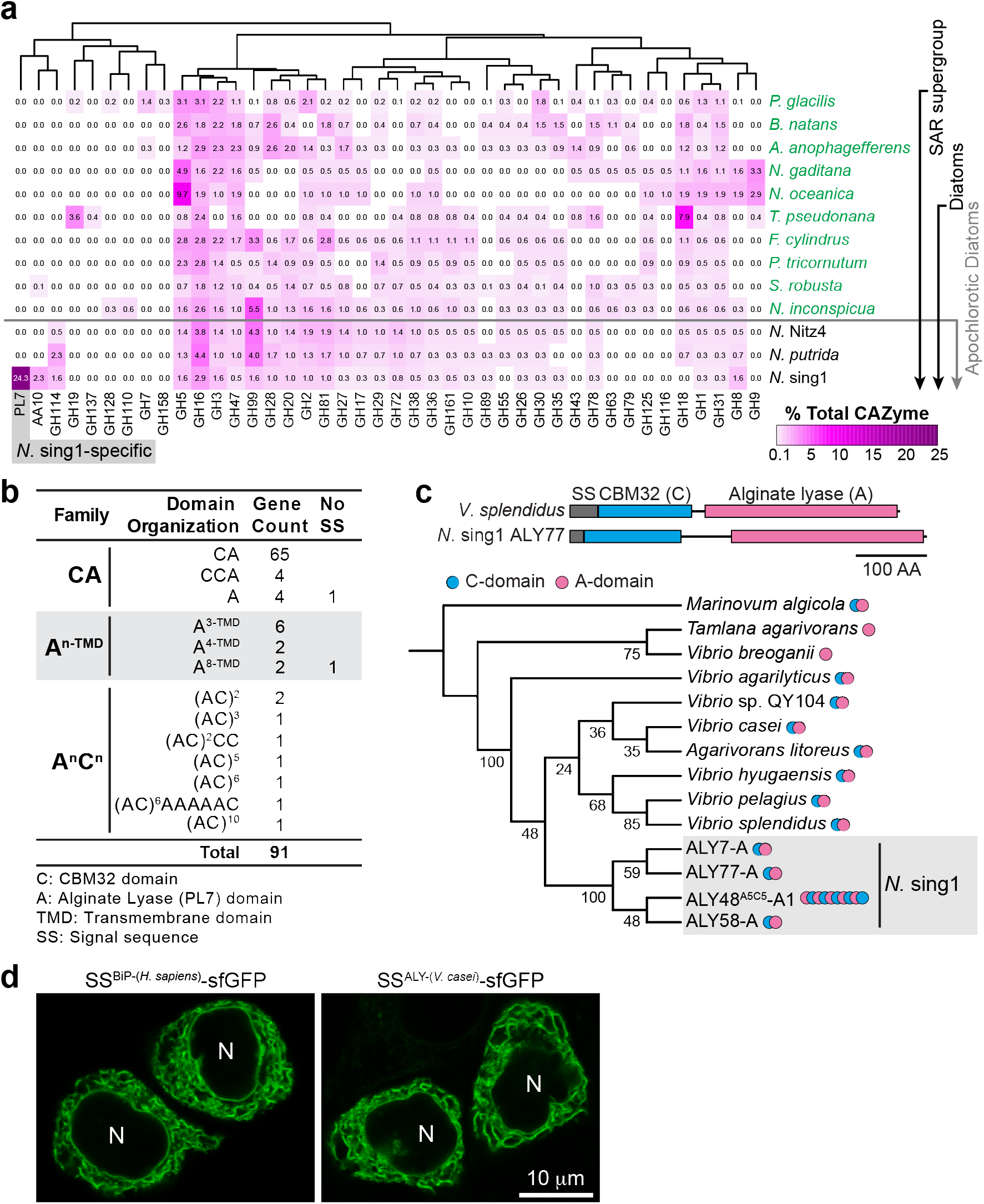
**Acquisition of alginate lyase (ALY) genes in *Nitzschia* sing1 by horizontal gene transfer (HGT). a**. Profile of selected carbohydrate-active enzymes (CAZymes) encoded in the genomes of select diatoms and other marine alga species (SAR supergroup). The heatmap displays the number of various CAZyme family genes as a percentage of the total CAZyme genes in each taxon (magenta scale, values indicated). The Polysaccharide Lyase 7 (PL7) family of *N.* sing1 is highlighted in gray. Related to Supplementary Fig. 2. **b**. Domain organization (N- to C- terminus) and number of predicted *N*. sing1 ALYs. ALYs are grouped into three families (CA, A^n-TMD^ and A^n^C^n^) based on overall domain organization. *n* denotes the number of PL7 alginate lyase (A) or CBM32 (C) domains in the predicted protein. TMD = transmembrane domain. The number of genes that do not encode a predicted signal peptide (no SS) is indicated. **c**. The schematic shows the domain arrangement of selected alginate lyase proteins from *Vibrio splendidus* and *N*. sing1 (ALY77). Maximum likelihood phylogenetic tree (bootstrap replicates = 1000) of *N*. sing1 ALYs and closely related bacteria ALYs. Note that *N*. sing1 ALYs form a monophyletic clade with a subset of *Vibrio* sp. alginate lyases. Related to Supplementary Fig. 3. **d**. The signal sequence from a *V. casei* alginate lyase directs a GFP fusion protein (SS^ALY-(*V.*^ *^casei^*^)^-sfGFP) into the endoplasmic reticulum (ER) of HeLa cells. The signal sequence from the human ER luminal protein BiP (SS^BiP-(*H.*^ *^sapiens)^*-sfGFP) serves as a positive control. N = nucleus. Scale bar = 10 μm.

*N*. sing1 occurs as an epiphyte on brown seaweeds and can catabolize alginate^32^. Thus, we focused on ALY genes as potential keys to understanding the basis for *N*. sing1’s adaptive habitat invasion. Alginate comprises up to 40 % of brown seaweed biomass^56^ and consists of linear chains of β-D-mannuronate (M) and its C5 epimer α-L-guluronate (G) linked by 1,4 glycosidic bonds. M and G residues can occur as Poly-M, Poly-G and alternating Poly-MG-enriched blocks^57^, which influences the physicochemical properties of the polymer^58^. Notably, Poly-G regions self-associate through divalent cation binding^59^ to form hydrogels, which play a structural role in the seaweed cell wall^60,61^.

To investigate the origin of ALY genes in the *N.* sing1 lineage, we performed protein sequence similarity searches (BLASTp) against the non-redundant (nr) protein database using all the *N*. sing1 ALY predicted proteins as query sequences. Each of the 91 ALYs returned bacterial PL7 family members from marine bacteria as top hits, with many of these also encoding N-terminal CBM32 domains. Phylogenetic analyses of these sequences show that *N*. sing1 ALYs form a monophyletic group with a subset of putative ALYs encoded by marine bacteria from the genus *Vibrio*. These findings strongly suggest that *N*. sing1 obtained an ancestral CBM32-containing ALY gene by horizontal gene transfer (HGT) from a *Vibrio* marine bacterium (Fig. 1c and Supplementary Fig. 3). The majority of *N*. sing1 ALYs encode predicted signal peptides, as do the nearest relative bacterial ALYs (Fig. 1b and Supplementary Table 4), suggesting that they are secreted. Bacterial secY- and eukaryotic Sec61-based secretory machineries have an ancestral relationship and act on hydrophobic signal sequences^62^. To assess the likely cellular fate of a bacterial alginate lyase in the eukaryotic cellular environment, we fused the signal sequence encoded by a *Vibrio casei* alginate lyase to sfGFP and expressed the fusion protein in mammalian HeLa cells. This protein is efficiently targeted to the endoplasmic reticulum (ER), presenting a fluorescent signal indistinguishable from that produced by sfGFP fused to the signal sequence from a *bona fide* ER-resident protein Binding Immunoglobulin Protein (BiP) (Fig. 1d). Thus, the ALY obtained by HGT from a marine bacterium is likely to have been immediately available for secretion by the ancestor of *N*. sing1.

### N. sing1 encodes three distinct alginate lyase gene families

The 91 *N*. sing1 ALY genes can be grouped into three families based on domain organization (Fig. 1b and Supplementary Table 4). The most abundant group (CA family) resembles the putative bacterial donor gene, encoding an N-terminal CBM32 domain (C-domain) followed by a single PL7 alginate lyase domain (A-domain). The two other families have undergone substantial changes in domain organization and appear to be the product of domain duplication, gain, loss and rearrangement. Members of the A^n-TMD^ family contain no C-domains but instead encode between 3 to 8 A-domains and a predicted C-terminal transmembrane domain. By contrast, members of the A^n^C^n^ family encode varying numbers of A- and C-domains arranged in repeat, with most encoding tandem AC repeats (Fig. 1b and Supplementary Table 4).

To investigate the relationship between these ALY genes, we constructed phylogenetic trees from the nucleotide sequences of all *N*. sing1 ALY A-domains (Fig. 2a and Supplementary Fig. 4). Importantly, A-domain sequences from the A^n-TMD^ and A^n^C^n^ families form a monophyletic sister clade to those from the CA family with 100 % bootstrap support. Moreover, each A-domain in these two families shares a 15-base pair (bp) coding sequence insertion at precisely the same position, suggesting that they diverged and expanded from a common ancestral gene (Fig. 2a, b). A small number of ALY genes lost or duplicated the C domain (‘CCA or A’ ALYs after their domain organization). These are nested as single genes within CA loci and occur throughout the CA phylogeny, suggesting that they arose independently multiple times (Fig. 2a and Supplementary Fig. 4) and represent a CA gene subfamily.

**Figure 2.**
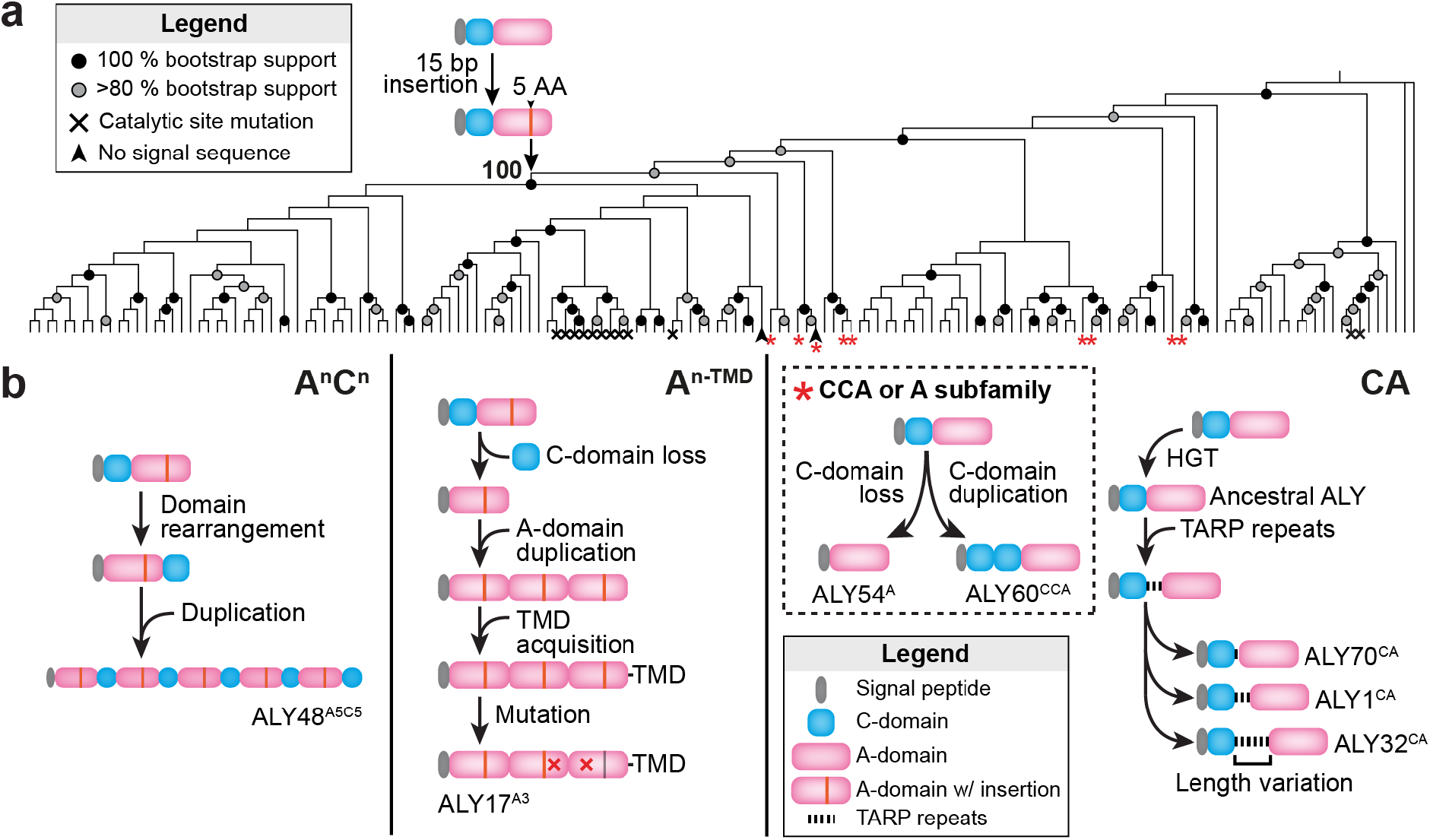
**Phylogeny, evolution and domain rearrangements of *N.* sing1 ALYs. a**. Maximum likelihood phylogenetic tree constructed from the nucleotide sequences of all *N*. sing1 ALY A-domains (bootstrap replicates = 1000). Note that domains from the A^n-TMD^ and A^n^C^n^ sequences form a monophyletic group (shaded in gray) with 100 % bootstrap support. These two families also share a 15 bp insertion, further indicating that they likely derived from a common ancestor. ALYs lacking a putative signal sequence are marked with a black arrowhead while sequences with mutations in conserved alginate lyase catalytic residues are marked with a black cross. ALYs belonging to the CCA or A subgroup in the CA family are marked with a red asterisk. Bootstrap support for nodes is summarized as indicated. Related Supplementary Fig. 4 and 8. **b**. Schematic diagrams illustrating the evolutionary genetic events leading to the different ALY families. Domain representations are identified in the legend. Note that the order of events is arbitrary. Specific ALYs are identified and shown for illustrative purposes.

To explore the mechanisms underlying ALY gene amplification and diversification, we next examined the structure of their genomic loci (Fig. 3a). The 91 ALY genes are distributed across 30 loci, 20 of which possess two or more genes in tandem. In most of these tandem loci (15/20), ALY genes occur in a unidirectional head-to-tail (5’ to 3’) orientation that conforms to duplication through unequal crossing over^63^. When such crossovers occur, intergenic regions are also duplicated (Fig. 3b). However, while coding regions can be retained through positive selection, these intergenic regions are expected to degenerate over time due to genetic drift. We next used nucleotide sequence dot plots^64^ to search for patterns of homology at duplicated ALY loci (Supplementary Fig. 5). We further developed a method to quantify these dot plots and graphically show levels of intergenic homology in a matrix plot (Fig. 3b and Supplementary Fig. 6). As expected, homologies are detected between all ALY coding regions at any given locus. In addition, some loci also display high levels of homology between consecutive intergenic regions (see loci 7, 8, 9, 19, 25, 26, 28 and 29) (Fig. 3b and Supplementary Fig. 6). Thus, these are likely to constitute the most recent duplication events. All these tandem loci are supported by long reads, suggesting that they are unlikely to be assembly artefacts related to repetitive DNA sequences (Supplementary Table 5).

**Figure 3.**
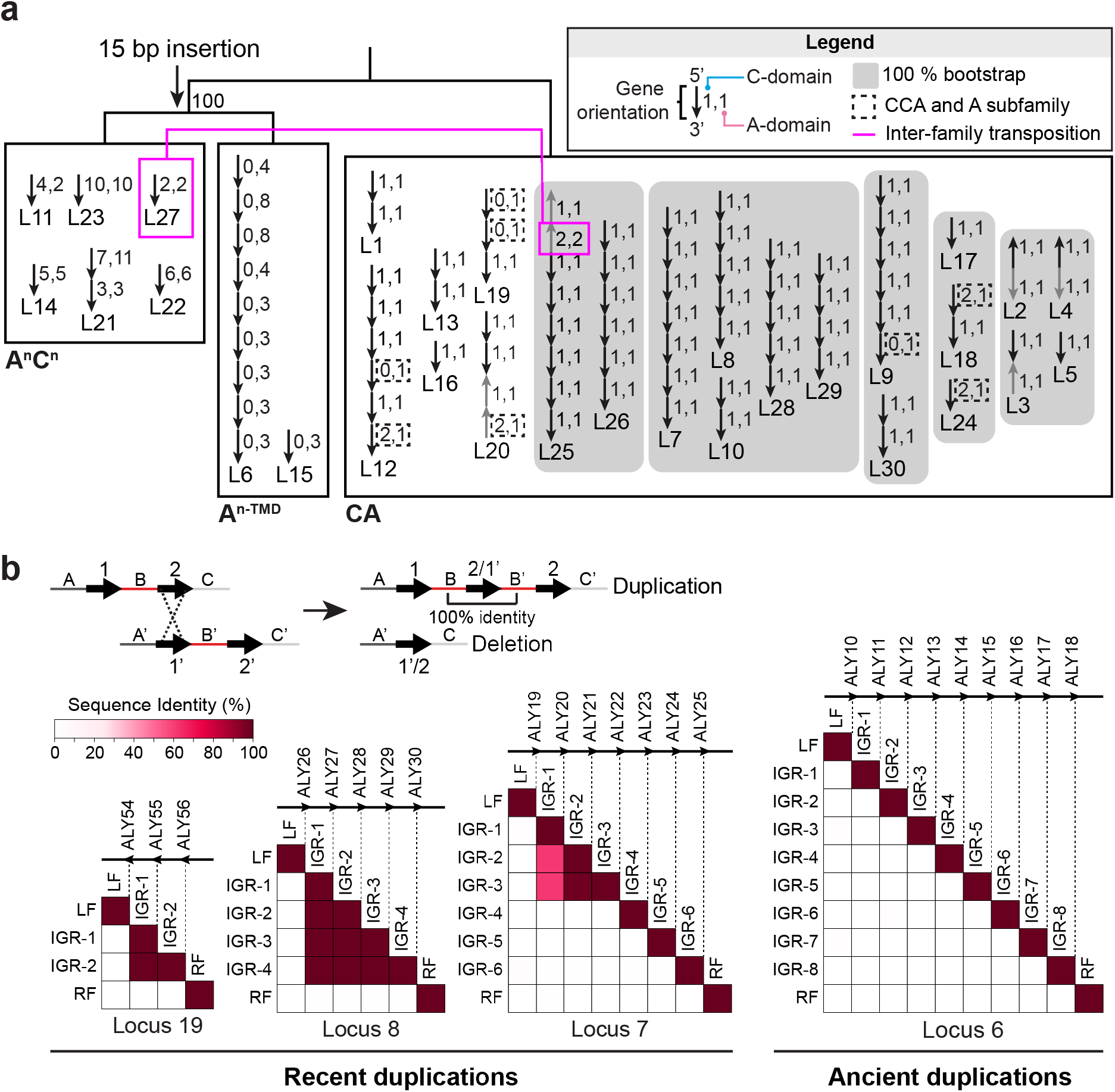
**ALY gene amplification through transposition and unequal crossing over. a**. A simplified A-domain phylogenetic tree (midpoint rooted) from Fig. 2a is shown along with the arrangement of genes at all 30 genomic loci. ALY families are labeled. Each arrow represents the orientation (5’ to 3’) of a single ALY gene. Genes oriented in opposite directions from the rest of the locus are colored in gray. Loci where most genes form a monophyletic clade with 100 % bootstrap support are grouped and are presumed to be related through transposition (gray background). The magenta lines link genes where A-domains form clades with 100 % bootstrap (support through at least one A-domain). These are likely to be related through transposition or gene conversion. The magenta line identifies a unique case where a transposition event appears to have introduced an A^n^C^n^ gene into a CA family locus to produce locus 25, which contains genes encoding ALYs from two different families. The numbers next to the arrows identify the number of C- and A-domains in the gene (e.g., 2,2 denotes two C- and two A-domain). **b**. Homology between intergenic sequences identifies recent tandem duplication events. The cartoon (top right) illustrates the result of unequal crossing over at a single locus containing two genes in tandem. Note that the intergenic region B is duplicated and has 100 % sequence identity immediately after the recombination event. The lower panels show self-comparison of intergenic regions (IGRs) from loci 6, 7, 8 and 19. The percent sequence identity is given for each pairwise comparison according to the heatmap. Recent duplication events (locus 7, 8 and 19) are evidenced by high levels of homology at consecutive intergenic regions. IGR: Intergenic Region; LF: Left Flank; RF: Right Flank. Related to Supplementary Fig. 5 and 6.

To investigate the role of transposition in ALY gene amplification, we correlated the phylogenetic relationships between ALY genes with the structure of their genomic loci (Fig. 3a). Specifically, we assign a relationship through transposition when all genes at two or more loci form monophyletic clades with strong (100 %) bootstrap support. These types of relationships are almost exclusively found within individual ALY families, suggesting that the three families evolved unique identities prior to amplification through transposition (Fig. 3a). Apart from the CA family loci containing CCA or A subfamily genes, only one locus contains genes from different families. Here, locus 25 consists of seven CA genes and one A^n^C^n^ gene (ALY67^A2C2^). Because the latter shows a close phylogenetic relationship to ALY82^A2C2^ encoded at locus 27, we conclude that the A^n^C^n^ gene at locus 25 arose through a transposition event. Only 3 of 20 tandem loci occur on contigs with lengths below 100 kb (Supplementary Fig. 7), indicating that assembly fragmentation is unlikely to account for the high number of ALY loci. Thus, transposition played a major role in expansion of the ALY gene family. Together, these findings document an initial ALY gene acquisition by HGT, diversification into three families through domain rearrangement and mutation, and ALY gene family expansion by tandem duplication and transposition.

### Enzymatic activity of N. sing1 alginate lyases

Bacteria generally initiate alginate catabolism by secreting endolytic PL7 family alginate lyases, which convert alginate polymers into short oligosaccharides. These are imported into the cell through various transporters before being converted into monomers by exolytic oligo-alginate lyases (OALs). Alginate monomers then enter central carbon metabolism to yield ATP and pyruvate^65^. Similarity searches failed to return any clear *N.* sing1 homologs of bacterial transporters or OAL enzymes (Supplementary Table 6), suggesting that these steps are carried out by distinct machineries.

We next examined the enzymatic activity of ALY gene products to determine how they contribute to alginate catabolism. PL7 alginate lyases possess well-defined catalytic residues^50,66^ that are largely conserved within the CA and A^n^C^n^ families (Fig. 2a). However, within the A^n-TMD^ family, many A-domains have substitutions at key catalytic residues (Fig. 2a and Supplementary Fig. 8). To examine ALY enzyme activity, we developed an assay where alginate gel liquefaction—which occurs with alginate degradation—is measured by fluid displacement upon vortexing (See Materials and Methods). A-domain representatives from each family were expressed in *E*. *coli* and crude extracts were examined for the ability to promote liquefaction. Despite some being found exclusively in inclusion bodies, all six CA family ALYs are active and display similar liquefying activities as compared to a positive control consisting of a commercially available endolytic alginate lyase. By contrast, A-domains from both A^n-TMD^ and A^n^C^n^ family members are inactive or display very low activity (Fig. 4a). We next directly measured endolytic enzyme activity through UV absorption by double bonds that form at non-reducing ends of oligosaccharide products. Attempts to purify soluble A-domains were met with limited success due to inclusion body formation. However, we were able to produce ALY7-A, ALY58-A, ALY77-A (CA family) and ALY48^A5C5^-A1 (A^n^C^n^ family) as soluble enzymes. Here, in keeping with the liquefaction assay, all CA family A-domains are enzymatically active, while the A^n^C^n^ family A-domain appears to be inactive (Fig. 4b).

**Figure 4.**
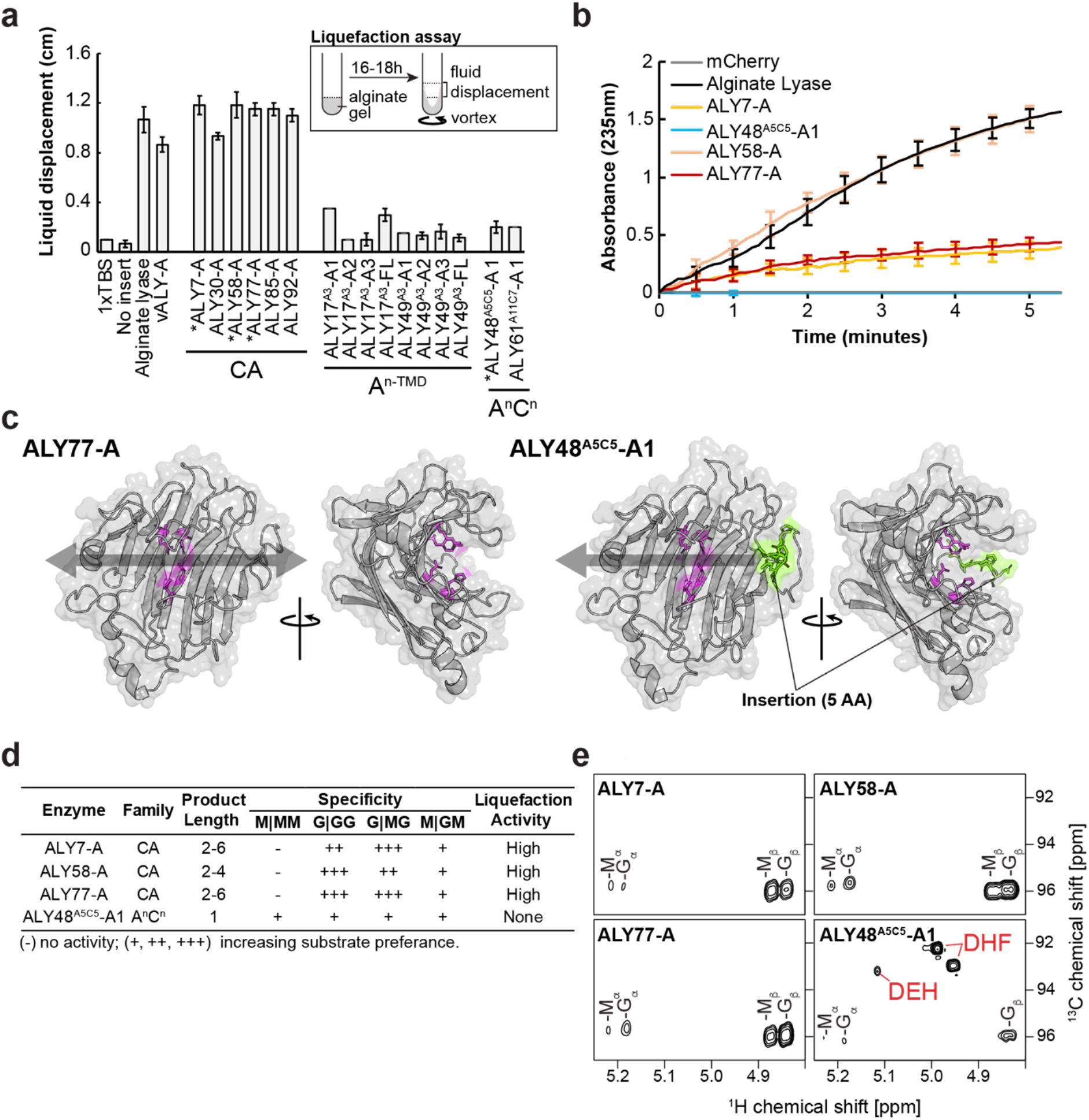
***N.* sing1 ALY enzyme activity and substrate specificity. a**. Alginate gel liquefaction assay using crude lysates from *E. coli* expressing the indicated proteins. Liquefaction is measured by the height of liquid displacement generated upon vortexing. Buffer alone (1×TBS) and a crude lysate from *E. coli* expressing an empty vector (No insert) serve as negative controls. Commercial endolytic alginate lyase (Alginate lyase) serves as a positive control. Note that all the CA family crude extracts promote liquefaction, while neither the A^n-TMD^ nor A^n^C^n^ family A-domains show substantial activity. Asterisks identify the proteins that could be purified in a soluble form. **b**. Alginate lyase activity of recombinant proteins as measured by an increase in absorbance at 235 nm. Enzymes are identified in the legend and standard deviation is shown. mCherry serves as a negative control, while a commercial endolytic ALY serves as a positive control. **c**. AlphaFold2 predictions of A-domains from representative CA (ALY77-A) and A^n^C^n^ (ALY48^A5C5^-A1) family proteins. Catalytic side chains are highlighted in magenta. Amino acid side chains encoded by the 15 bp insertion are colored in green. The opaque black arrows show how an alginate polymer might bind to the enzymes. Related to Supplementary Fig. 9. **d**. The table summarizes the enzymatic activity of the indicated recombinant proteins with deduced enzyme specificity and reaction products identified. **e**. Reaction products produced by ALY enzymes acting on seaweed alginate. The panels show a region of the NMR spectrum (^1^H-^13^C HSQC) where alginate monomers DEH (4-Deoxy-L-erythro-5-hexoseulose uronate) and DHF (two epimers of 4-deoxy-D-manno-hexulofuranosidonate) occur (labelled in red). The other annotated C/H-1 signals correspond to: Mα, α-D-mannuronate reducing end; Mβ, β-D-mannuronate reducing end; Gα, α-L-guluronate reducing end; Gβ, β-L-guluronate reducing end. Related to Supplementary Fig. 10.

To investigate how the 15-bp insertion in the A-domain of ALY48^A5C5^-A1 might impact its enzyme activity, we predicted its structure using AlphaFold2^67^ and compared it to the active endolytic CA enzyme, ALY77-A. The predicted structure of both proteins closely resembles the β-jelly roll fold seen in PL7 alginate lyase crystal structures^50^ (Fig. 4c and Supplementary Fig. 9). Interestingly, in ALY48^A5C5^-A1, the 5 amino acids encoded by the 15-bp insertion forms a surface loop that appears to occlude the alginate-binding groove on one end to form a binding pocket. This could promote alginate end-binding and convert the enzyme from an endolytic to exolytic mode of action. Such an occlusion has previously been correlated with the emergence of exolytic activity in a PL7 family alginate lyase (AlyA5) from the bacterium *Zobellia galactanivorans*^68^.

To determine the specificity of endolytic CA family A-domains and the possibility that ALY48^A5C5^-A1 is an exolytic alginate lyase, we next employed NMR to directly observe reaction products^69^ (Fig. 4d, e and Supplementary Fig. 10). Substrates examined include seaweed alginate and defined polymers composed of Poly-M, Poly- MG and Poly-G tracts. These data confirm that all three CA family A-domains are endolytic enzymes that accumulate oligosaccharides between two and six residues in length. In addition, they are inactive on Poly-M and generally show distinct preferences for cleavage at G|GG and G|MG sites. None of these enzymes produced monomers, indicating that they are exclusively endolytic. By contrast, the ALY48^A5C5^-A1 domain displays an exolytic activity on all alginate substrates and accumulates DEH (4-Deoxy-L-erythro-5-hexoseulose uronate) and DHF (4-deoxy-D-mannohexulofuranosidonate), which arise spontaneously from the primary monomer product (4,5-unsaturated uronic acid). Together, these findings indicate that CA family members have retained the ancestral endolytic function, while the A^n^C^n^ A-domain acquired an exolytic mode of action (Fig. 4c, d and Supplementary Fig. 10).

### A role for vacuoles and endomembrane trafficking in alginate uptake

To better understand the cell biology of *N*. sing1 alginate catabolism, we imaged *N*. sing1 cells grown in seawater medium supplemented with alginate or dextrose as sole carbon sources. Alginate forms hydrogels in seawater that can be seen as amorphous aggregates with brightfield microscopy (Fig. 5a). Over time, *N*. sing1 diatoms begin to degrade alginate, as evidenced by the increase in absorbance at 235 nm. The medium also turns yellow and develops a broad absorbance shoulder that tails off at around 500 nm (Fig. 5b). In addition, alginate hydrogels also start fluorescing when viewed under epifluorescence (Fig. 5a). Interestingly, vacuoles of diatoms grown on alginate display a related fluorescence, while those grown on dextrose do not (Fig. 5c). The emission spectra of these vacuoles and the alginate seawater medium from a saturated *N.* sing1 culture overlap (Fig. 5e), suggesting that the vacuole-derived fluorescence is indeed due to uptake of alginate-related fluorescent moieties. To further examine this phenomenon, we employed the vacuole dye CMAC (7-amino-4-chloromethylcoumarin)^70^ and found that vacuoles in alginate- grown cells are substantially larger and more abundant than those observed in dextrose-grown cells (Fig. 5c, d and Supplementary Fig. 11a). Together, these data suggest that products of alginate breakdown are taken up by endocytosis and processed within vacuoles. When *Vibrio* sp. bacteria are grown on the same alginate medium, absorbance at 235 nm is detected. However, no additional absorbance (Fig. 5b) nor medium yellowing is observed (Supplementary Fig. 11b). Thus, these aspects of alginate metabolism appear to be *N.* sing1-specific.

**Figure 5.**
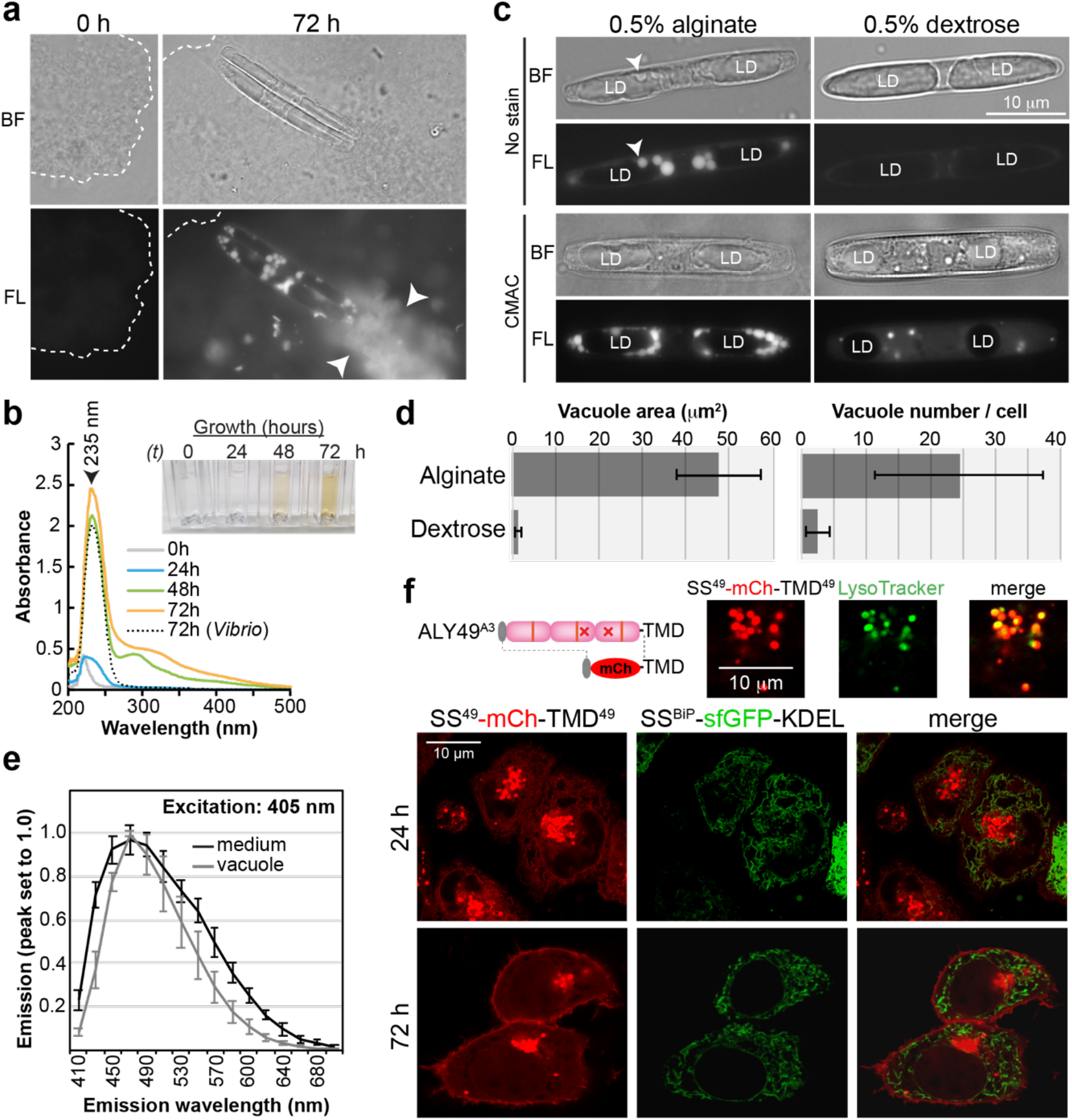
**Cell biology of diatom alginate metabolism. a**. Alginate hydrogels in seawater medium are seen as amorphous aggregates by brightfield microscopy (BF, outlined). Note that they begin to fluoresce as *N*. sing1 diatoms grow (72 h, arrowheads). Scale bar = 10 μm. **b**. Absorbance spectra of alginate culture medium from an *N*. sing1 culture and a *Vibrio* sp. culture at different time points. The absorbance peak at 235 nm corresponds to non-reducing ends produced by alginate lyase activity. Note the absorbance shoulder (270–450 nm) that increases over time with *N*. sing1 cultures, but not *Vibrio* sp. cultures. The inset shows the appearance of diatom culture media at various time points. Related to Supplementary Fig. 11b. **c**. Vacuoles in *N*. sing1 (arrowheads) fluoresce when grown on alginate but not when grown on dextrose (LD: lipid droplets). CMAC staining reveals the proliferation of vacuoles in alginate as compared to dextrose cultures. **d**. Quantification of vacuole size and number in cells grown on 0.5 % alginate or 0.5 % dextrose (n = 10). Related to Supplementary Fig. 11a. **e**. Emission spectra (excitation: 405 nm) of medium (black) and *N*. sing1 vacuoles (gray) after 72 h growth. **f**. Localization of mCherry fusion protein with the signal peptide (SS) and transmembrane domain (TMD) from ALY49^A3^ in HeLa cells (SS^49^- mCh-TMD^49^). The perinuclear bodies are defined as lysosomes by colocalization with LysoTracker (upper panels). sfGFP fused with a signal sequence from the human endoplasmic reticulum (ER) protein BiP and the ER retention signal (KDEL) serves as a marker for the ER (SS^BiP^-sfGFP-KDEL). Note that SS^49^-mCh-TMD^49^ localization shifts from the ER and lysosomes at 24 h to the plasma membrane and lysosomes at 72 h. Scale bar = 10 μm.

A^n-TMD^ family members possess mutations in catalytic domains (Supplementary Fig. 9) and appear to be enzymatically inactive (Fig. 4a). These all possess a predicted N-terminal signal sequence and a family-specific C-terminal transmembrane domain (TMD). This configuration is expected to result in cell surface proteins anchored to the plasma membrane with A-domains projecting into the extracellular milieu. Based on this, we hypothesize that they function as alginate import receptors. As attempts to transform *N*. sing1 to express an A^n-TMD^ fluorescent fusion protein were unsuccessful, we turned to heterologous expression in a mammalian cell line. A full-length mCherry- ALY49^A3^ fusion protein did not appreciably accumulate. However, we were able to visualize an mCherry fusion protein encoding the N-terminal signal sequence and C-terminal transmembrane domain from ALY49^A3^ (Fig. 5f). At early time points after transfection (24 h), the fusion protein is detected in both the ER and lysosome. This indicates a signal peptide-dependent insertion into the ER, followed by endomembrane trafficking to the lysosome. At later time points (72 h), the fusion protein localizes to the cell surface and lysosome, but not in the ER (as expected of transient transfections). This pattern of steady-state localization suggests cycling between the plasma membrane and lysosome, which is consistent with the expected behavior of a eukaryotic import receptor.

### Disordered repetitive domains evolved in the N. sing1 lineage and bind to alginate

Data presented thus far document the evolution of ALY protein novelties in enzymatic activity (Fig. 4c-e) and protein trafficking (Fig. 5f). We next examined low-complexity sequences that occur between the two domains of CA family proteins (Fig. 2a). These are predicted to be disordered (Supplementary Fig. 12a) and are enriched for tetrapeptide sequences with a T-(A/P/S/V)-R-P consensus motif (‘TARP’ repeats). Importantly, these repeats are not observed in bacterial alginate lyases, indicating that they evolved independently in the *N*. sing1 lineage. Protein sequence alignments show that the length of these TARP-containing regions tends to vary precisely in the number of TARP repeats (Fig. 6a and Supplementary Fig. 12b). These regions also have a net positive charge through arginine that could bind to negatively charged polysaccharides like alginate. To explore this idea, we produced mCherry fusion proteins to a diverse group of TARP repeat regions (Supplementary Fig. 13a), along with a C-domain from the CA family gene ALY77 (ALY77-C). ALY77-C-mCherry binds to calcium alginate hydrogels as evidenced by fluorescence imaging (Fig. 6b and Supplementary Fig. 13b), a pelleting assay (Supplementary Fig. 14a) and fluorescence recovery after photobleaching (FRAP) (Supplementary Fig. 14b). TARP repeat regions also bind to alginate gels; however, they produce fluorescent puncta distinct from the uniform binding pattern observed with the C-domain (Fig. 6b and Supplementary Fig. 13b). In general, naturally occurring TARP sequences with more repeats produce brighter fluorescent puncta (Supplementary Fig. 13b). To exclude the influence of other sequences found in these regions, we next produced mCherry fusions with varying lengths of consecutive TARP repeats (3, 6, 9, 12). Fluorescence microscopy shows that alginate binding by these sequences is indeed cooperative, with those containing more TARP repeats producing brighter fluorescence signal (Fig. 6b).

**Figure 6.**
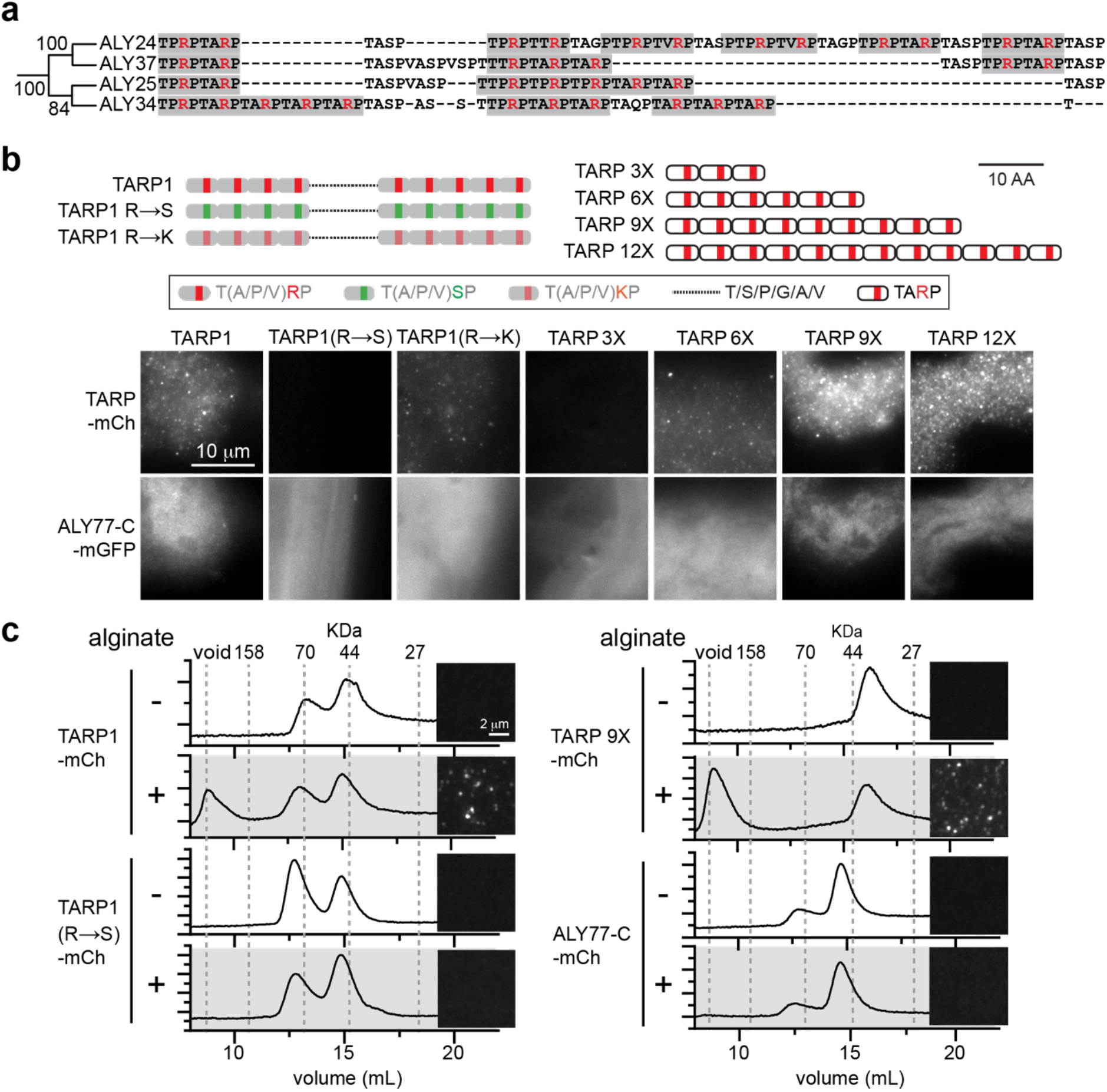
**TARP repeat sequences bind to alginate. a**. Closely related CA proteins show variation in TARP repeat length. TARP repeats are highlighted in gray and arginine (R) residues are colored red. Related to Supplementary Fig. 12b. **b**. The cartoons depict TARP repeat variants used in this study. TARP1 is a naturally occurring sequence found between the C- and A-domains of ALY1. R®S and R®K denote TARP1 with arginine substitutions to serine and lysine, respectively. TARP 3/6/9/12X are synthetic TARP repeat length variants. TARP repeat sequences fused to a mCherry protein bind calcium alginate hydrogels to produce a punctate pattern of fluorescence distinct from the uniform signal produced by CBM32^ALY77^ (C^ALY77^). R®S substitution abolishes binding while R®K substitution diminishes binding. Length variants show that TARP repeats promote binding in a cooperative manner. Scale bar = 10 μm. Related to Supplementary Fig. 13 and 14. **c**. TARP repeats bind to soluble alginate. Gel filtration chromatograms of TARP variants and C^ALY77^ in the presence (+) or absence (-) of alginate. A shift to the void volume is observed in both TARP1-mCh and TARP 9X-mCh when alginate is added, indicating the formation of a macromolecular complex. This shift is not observed with C^ALY77^ or the TARP1 R®S variant. Images next to each chromatogram show the same samples imaged by epifluorescence. Note that fluorescent puncta are only observed in samples that display a void volume shift in the chromatogram. Scale bar = 2 μm.

To investigate the role of arginine in TARP repeats, we chose the ALY1 TARP domain (TARP1), which contains nine TARP repeats, and produced variants that substitute arginine with either serine or lysine. Substitutions with serine (R→S) abolish alginate binding, while substitutions with positively charged lysine (R→K) diminish binding (Fig. 6b). These data indicate that electrostatic interactions promote alginate binding by TARP repeats. To investigate whether TARP and C-domains also bind to uncomplexed alginate, we incubated the recombinant fusion proteins with alginate in the absence of calcium. Fluorescence microscopy shows that TARP proteins form complexes with soluble alginate to produce a fine punctate signal. By contrast, the C-domain and TARP1 R→S variant do not produce any detectable signal (Fig. 6d). These observations are corroborated by gel filtration, where a shift in the TARP1 elution peak to the void volume indicates formation of a high molecular weight complex between TARP1 and alginate. This shift is abolished with the TARP1 R→S variant and does not occur with the C-domain (Fig. 6d). Together, these data show that C-domains bind to structural moieties associated with the calcium alginate hydrogel, while TARP repeats appear to bind uncomplexed alginate through arginine-based electrostatic interactions.

## Discussion

Apochlorotic diatoms lost photosynthesis to become free-living secondary heterotrophs. Here, we account for their acquisition of alginate catabolism and adaptive invasion of brown seaweed habitats. The ancestor of *N*. sing1 appears to have acquired an alginate lyase gene by HGT from a *Vibrio* marine bacterium (Fig. 1). From this basis, the physiological capacity to degrade alginate evolved through gene amplification and neofunctionalization (Fig. 2). Tandem duplication and transposition underlie amplification (Fig. 3), while domain rearrangement, gain, loss, mutation, and *de novo* sequence evolution led to the advent of new protein activities. These include alginate lyases with exolytic enzyme activity (Fig. 4), putative alginate import receptors (Fig. 5) and low-complexity alginate-binding sequences (Fig. 6). Thus, while HGT from a primary heterotroph provided the foundation for alginate catabolism, evolving the full capacity required a substantial amount of lineage-specific genetic innovation.

The ancestral endolytic function of the PL7 family alginate lyases is retained in *N*. sing1 CA family ALYs (Fig. 3). However, alginate catabolism also requires mechanisms for the import of alginate oligosaccharides and their digestion into monomers. Marine bacterial alginate degradation systems are well- characterized^48,50,65^: In *Vibrio* sp., extracellular alginate oligosaccharides produced by endolytic alginate lyases^48,50^ are imported through transporters^65^ and subsequently converted to monomers by cytoplasmic OAL enzymes. *N*. sing1 lacks bacteria-like transporters and OALs (Supplementary Table 6). Instead, monomer production can be accounted for by the emergence of exolytic activity in the A^n^C^n^ family A-domains (Fig. 4d, e and Supplementary Fig. 10), while alginate import appears to depend on the endomembrane system (Fig. 5c), with A^n-TMD^ family members potentially acting as import receptors (Fig. 5f). Alginate-binding TARP repeats represent an additional form of innovation that appears to be unique to the *N*. sing1 lineage.

HGT can drive eukaryotic evolution when an acquired gene promotes access to a new ecological niche or lifestyle. Such acquisitions have been linked with adaptation to environmental extremes, parasitism, and nutrient acquisition (reviewed in^71–74^). Genes acquired by HGT appear to frequently undergo duplication^75–81^. However, relatively few studies have performed functional assays to document new protein activities in duplication-derived paralogs^82–85^. In two such studies, duplication-derived paralogs evolved new activities related to nutrient acquisition^83,84^, as documented here for ALY genes.

Most ALY gene tandem duplicates are arranged in a head-to-tail orientation, suggesting that they expanded through unequal crossing over during homologous recombination. High levels of intergenic sequence homology at certain ALY loci provide definitive evidence for this conclusion (Fig. 3b, Supplementary Fig. 5 and 6) and indicate that the evolution of alginate catabolism is likely to be ongoing. This mechanism can account for gene duplication as well as the domain duplications required for the emergence of A^n-TMD^ and A^n^C^n^ families (Fig. 2 and 3). We conclude that unequal crossing over was a major source of genetic variation underlying the evolution of alginate catabolism. Unequal crossing over may also account for the expansion of low-complexity TARP repeat sequences. However, in this case, an origin through other mechanisms such as errors in DNA replication cannot be excluded.

Expansion of tandem gene duplicates has been associated with the evolution of environmental sensing in plants^86^, and adaptation to benthic marine habitats in diatoms^40^ and shrimp^87^. To the best of our knowledge, intergenic homology has not been employed to identify recent expansion events as shown here for ALY loci (Fig. 3b). The systematic application of this approach has the potential to identify actively expanding loci related to ongoing adaptive change. Interestingly, lab-based evolution experiments in the diatoms *Seminavis robusta* and *Phaeodactylum tricornutum* documented high rates of mitotic homologous recombination^88^. This suggests that duplications may frequently arise through unequal crossing over in vegetative populations. These findings raise a scenario where much of the genetic variation that led to the evolution of alginate catabolism in the *N.* sing1 lineage had an asexual origin. Lab-based evolution experiments with *N.* sing1 will help to address this possibility.

The CA family expanded to comprise 73 members and the majority of these retain signal sequences and catalytic residues (Fig. 2). Gene dosage effects and diversification of enzyme specificity towards different alginate linkages are likely to provide a selective advantage leading to CA gene expansion and maintenance. Interestingly, in closely related *Vibrio* strains, HGT leading to high alginate lyase copy number is correlated with increasing enzyme activity and efficiency of alginate utilization^89^. Thus, gene dosage effects appear to have played a role in the adaptive evolution of alginate catabolism in both bacteria and eukaryotes. The C-domain was acquired from the founding bacterial ALY gene, while TARP repeats appear to have arisen in the *N.* sing1 lineage. Both bind to *Sargassum* cell walls (Supplementary Fig. 14c) but interact with alginate through distinct mechanisms (Fig. 6c). Future work can examine whether these domains cooperate and how they influence alginate processing by A-domains.

A^n-TMD^ family members acquired a C-terminal transmembrane domain (Fig. 2a and 4a) and lost enzymatic activity (Fig. 4a) through mutation (Fig. 2b). Heterologous expression (Fig. 5e) indicates that A^n-TMD^ family members are likely to be tethered to the cell surface to present A-domains to the extracellular space. Growth of *N*. sing1 on alginate is accompanied by the formation of alginate-related fluorescent moieties that enter vacuoles (Fig. 5a). Moreover, vacuoles become enlarged and increase in numbers when diatoms are grown on alginate as compared to dextrose (Fig. 5c). Thus, while the nature of fluorescent alginate-related moieties remains to be determined, endocytosis and vacuoles appear to provide for alginate internalization. A^n^C^n^ family members have signal sequences for entry into the secretory pathway (Fig. 1b) and can convert alginate oligosaccharides into monomers (Fig. 4e). However, whether they are secreted outside the cell or undergo endomembrane transport to obtain residency in the vacuole remains to be determined. Other open questions concern how alginate-derived sugars are transported from the vacuole lumen to the cytoplasm, as well as how they enter central carbon metabolism. Additional work and the development of techniques for genetic transformation and gene editing in *N.* sing1 will help to address these questions.

Marine fungi comprise the other group of eukaryotes that are known to catabolize alginate^90^. The genome sequence and proteomic analysis of one such fungus, *Paradendryphiella salina,* identified three secreted PL7 family alginate lyases that are induced by the presence of alginate^91^. These data support the idea that marine fungi and apochlorotic diatoms initiate alginate catabolism through secreted endolytic alginate lyases. Interestingly, another marine fungus, *Asteromyces cruciatus,* encodes a fungus-specific transporter, Ac_DHT1, which confers alginate monomer import upon heterologous expression in yeast^92^. *N.* sing1 does not encode Ac_DHT1 homologs (Supplementary Table 6). Thus, marine fungi and diatoms appear to be distinct in their mechanism of alginate import.

Diatoms are likely to possess features that predispose them to a successful transition to obligate heterotrophy. These include high levels of intraspecific genetic variation^37,40,93^, a predisposition towards rapid evolution^37,39,40,88^, mixotrophy^94–97^, diverse metabolic networks^98–101^, and substantial force generation through gliding motility^32^. *N*. sing1 and phylogenetically distinct relatives catabolize alginate, while *N*. *putrida* does not^32^. Moreover, *N*. *putrida* and *N*. Nitz4 do not encode ALY genes (Fig. 1a). Thus, current data point to a high degree of ecophysiological diversity in the apochlorotic lineage. More environmental sampling and genome sequencing will be required to better understand the biodiversity and heterotrophic strategies of these diatoms. Going forward, apochlorotic diatoms will make good models to study the genetics and physiology of adaptive radiation while also expanding our understanding of nutrient cycling at the intertidal zone.

## Materials & Methods

### Diatom culture conditions and nucleic acid extraction

*Nitzschia* sing1-1 diatoms were cultured on synthetic seawater (SSW) medium as previously described^32^. For genomic DNA (gDNA) and total RNA extraction, diatoms were grown in liquid SSW medium supplemented with 0.5 % (w/v) dextrose (Sigma-Aldrich, D9434) at 30°C for 3 days. For cell imaging experiments (Fig. 5a-e), diatoms were grown in SSW medium supplemented with either 0.5 % (w/v) dextrose or medium-viscosity sodium alginate (Sigma-Aldrich, A2033). *N.* sing1 gDNA was extracted using the MasterPure™ Yeast DNA Purification Kit (Lucigen, MPY80200). Total RNA was extracted with TRIzol™ (Invitrogen, 15596018), treated with DNase I (Roche, 04716728001) at 37°C for 20 minutes and purified by Phenol/Chloroform extraction.

### Genome sequencing and assembly

*N*. sing1 gDNA was sequenced using the Pacific Biosciences (PacBio) single-molecule real-time sequencing (SMRT) technology (Sequel II system, Continuous Long Read (CLR) mode) (NovogeneAIT Genomics, Singapore) for long-reads and BGI’s cPAS and DNA nanoballs technology (BGISEQ-500, DNB-seq) (BGI Tech Solutions (Hong Kong) Co., Ltd.) for short-reads. PacBio long-read sequencing yielded 7.75 Gb (608,716 subreads, mean read length = 12.5 kb) after filtering (SMRT Link parameters: minLength 0, minReadScore 0.8) and partitioning. Paired-end short-read sequencing yielded 4.24 Gb (28,322,876 reads, read length = 150 bp). Adapter and low-quality short-read sequences were filtered using the SOAPnuke^102^ software (SOAPnuke parameters: -n 0.01 -l 10 -A 0.25 -Q 2 -G -minLen 150).

PacBio sequence long-reads were assembled using HGAP4 (Hierarchical Genome Assembly Process 4)^103^ from SMRT Tools (SMRT Link v10.1.0 package), resulting in 80 contigs with a total length of 40.4 Mb. This initial assembly was subjected to two rounds of correction with Pilon (v1.24)^104^ based on genomic BGISEQ short-reads. BLAST comparisons^105^ against mitochondrial proteins from *Nitzschia alba* (GenBank accession: NC_037729.1)^106^ revealed two contigs that most likely correspond to mitochondria DNA. These two contigs, which also have a much lower GC content (29 %) than other contigs (45 %), were removed from the final assembly. The final genome assembly consists of 78 contigs with a total size of 40.3 Mb (Supplementary Table 1).

To assess the quality of the genome assembly, BUSCO (Benchmarking Universal Single-Copy Orthologs, v5.5.0)^107^ assessments were performed using the eukaryotic (eukaryote_odb10) and stramenopiles lineage datasets (stramenopiles_odb10). Genome assembly statistics were also computed with QUAST (v5.2.0)^108^ (Supplementary Table 1). *K*-mers were counted using the Jellyfish software (v2.3.1) and genome heterozygosity was visualized using GenomeScope (v1.0.0) (Supplementary Fig. 1). Analysis of transposable and other repeat elements was performed as described in Traeger *et al*. (2013)^109^. Briefly, repeats in the *N.* sing1 genome were identified using RepeatMasker (v4.1.5; rmblastn engine v2.14.0^110^) with a *de novo* repeat library generated from RepeatModeler (v2.0.5)^111^. Putative telomeric repeats (sequence TTAGGG) were identified using a custom Perl script (Supplementary Table 2).

BLAST analysis on one of the two putative mitochondrial contigs showed that they contain all the proteins predicted in the *N. alba* mitochondrial genome^106^. These contigs also have a similar size of 37 kb (compared to 36.2 kb of *N*. *alba*), further suggesting that they represent the mitochondrial genome of *N*. sing1. Contigs representing plastid DNA were not present in the PacBio assembly. However, a BLAST search of a BGISeq-only assembly generated with SPAdes (v3.14.0)^112^ against the *Nitzschia* Nitz4 plastid (GenBank accession: MG273660.1)^25^ identified three contigs with a total size of 59 kb and a GC content of 25 %, which most likely represent the plastid genome.

### Gene prediction and functional annotation

Genome annotation of protein-coding genes for the nuclear genome was performed using the MAKER (v3.01.03)^113^ and BRAKER2 pipelines (v2.1.2)^114^. Predicted proteins from the diatoms *Fistulifera solaris* (GenBank accession: GCA_002217885.1)^115^, *Pseudo-nitzschia multistriata* (GenBank accession: GCA_900660405.1)^116^, *Phaeodactylum tricornutum* (GenBank accession: GCF_000150955.2)^39^, *Fragilariopsis cylindrus* (GenBank accession: GCA_001750085.1)^37^, *Chaetoceros tenuissimus* (GenBank accession: GCA_021927905.1)^117^, Fragilaria crotonensis (GenBank accession: GCA_022925895.1)^118^, and Nitzschia inconspicua (GenBank accession: GCA_019154785.2)^38^ were used as inputs for both pipelines. For MAKER, the *N.* sing1 transcriptome assembled from two RNA-seq datasets with Trinity (v.2.4.0)^119^ was also included.

Two independent annotations were generated using BRAKER2. The first set of annotations (BRAKER2 parameter: --etpmode) was generated based on the diatom protein datasets used as input and the BAM files of the two RNA-seq datasets mapped to the *N*. sing1 genome assembly with Hisat2 (v2.2.1)^120^. The second annotation set merges the results from two BRAKER2 runs (using TSEBRA v1.0.3)^121^, which used either the protein or RNA-seq datasets as input. To derive at a final annotation set, all three gene model predictions from MAKER and BRAKER2 were combined using a custom Perl script. To assess the quality of the protein annotation, BUSCO (v5.5.0) analysis was performed using datasets from the eukaryote (eukaryote_odb10) and stramenopiles lineage (stramenopiles_odb10) (Supplementary Table 1). Putative mitochondria and plastid contigs were annotated using Prokka (v1.10)^122^.

Functional annotation of *N*. sing1 protein-coding genes was performed by searching for conserved protein domains using hmmscan (Parameters: --domtblout mode, domain i-evalue < 1.0E^-03^) from HMMER (v3.3.2) against the Pfam protein database^123,124^. The signal sequence for each protein was predicted using a combination of Phobius (webserver)^125^ and SignalP6.0^126^ (Supplementary Table 4). Annotation of carbohydrate-active enzymes (CAZymes) was also performed with hmmscan (--domtblout mode, domain i-evalue < 1.0E^-10^) using the dbCAN2 HMM database (HMMdb-V11)^127^. Proteins were assigned a CAZy family using the top EC hit with the lowest e-value. Heatmaps depicting the proportion of CAZymes in each genome were clustered using the Bray-Curtis dissimilarity measure. Scores for each EC/CBM class were calculated by normalizing the count against the total number of CAZyme within each species (Fig. 1a and Supplementary Fig. 2).

### N. sing1 alginate lyase nomenclature

ALY genes are numbered according to their position on contigs ordered from the largest to smallest. For proteins with single alginate lyase and/or CBM32 domains, they are referred to as ‘A’ or ‘C’, respectively. For example, the A domain of ALY1 is referred to as ALY1-A. In cases where proteins encode more than one A- or C-domain, the domains are numbered by type according to their position from N- to C-terminus. For example, the third A domain of the A^n^C^n^ family protein ALY48^A5C5^ is referred to as ALY48^A5C5^-A3. TARP repeat sequences are numbered corresponding to the ALY gene from which they originate. For example, the TARP repeat from ALY1 is designated TARP1. In cases where TARP repeats are identical between ALY genes, they are identified by the first ALY gene in which they occur.

### Phylogenetic and sequence analysis

To identify the closest relatives to *N*. sing1 ALYs, BLASTp searches against the NCBI non-redundant database were performed with all 91 *N.* sing1 ALY sequences (BLASTp parameters: max target sequences 500, e-value threshold 0.05, word size 5, matrix: BLOSUM62, gap extension cost: 11, gap opening cost, 1, all other parameters default). The BLASTp output for each ALY query sequence was merged into a single non-redundant dataset consisting of 27,320 sequences. These were then binned into 10 groups by bit score with an interval of 50. 25 sequences from each group were randomly sampled for phylogenetic tree construction. In this manner, we analyzed representative sequences from the full BLASTp output. To avoid poor alignment between sequences due to differences in domain organization, phylogenetic analysis was performed using A-domains only. For proteins with multiple A-domains, each domain was included as separate sequences. Multiple sequence alignment was prepared with MAFFT (v7.490)^128^ using the L-INS-I iterative refinement method and maximum likelihood phylogenetic trees were constructed using IQ-TREE2 (multicore v2.2.6) (Parameters: -b 1000 (bootstrap replicates))^129,130^ (Fig. 1c and Supplementary Fig. 3).

To investigate the relationship between *N*. sing1 ALY genes, nucleotide sequences from all A-domains were compared and used to construct maximum likelihood phylogenetic trees (Fig. 2a and Supplementary Fig. 4). For proteins with multiple A-domains, each domain was included as separate sequences in the respective trees. Multiple sequence alignment and phylogenetic analysis were performed as above. TARP repeat sequences (Fig. 6, Supplementary Fig. 12 and 13) were defined in all CA family proteins using the C- and A-domains as boundaries.

Multiple sequence alignment of TARP repeat sequences was performed with MAFFT (v7.490) using the E-INS-I iterative refinement method and maximum likelihood phylogenetic trees were constructed using IQ-TREE2 (multicore v2.2.6) (Parameters: -b 1000 (bootstrap replicates)^129,130^ (Fig. 6a and Supplementary Fig. 12b). For the alignment in Supplementary Figure 12b, sequences were manually aligned to show TARP repeat expansions. Disorder in TARP repeat regions were predicted using IUPred3^131^ (Supplementary Fig. 12a). Visualization of sequence alignments around key A-domain catalytic regions was made using ESPript3.0^132^ (Supplementary Fig. 8), while phylogenetic trees were visualized using iTOL (v6.9)^133^.

### Protein structural prediction

Structural predictions for A-domains were made using AlphaFold2 (ColabFold v1.5.5; AlphaFold2 using MMseqs2) with default parameters^134^ (Fig. 4c and Supplementary Fig. 10). For ALY48^A5C5^, only the first A-domain (ALY48^A5C5^-A1) as annotated by hmmscan was used for the prediction. The A-domain of AlyB from *V. splendidus* was obtained from PDB (PDB ID: 5ZU5)^53^. All visualizations were made and captured with PyMOL (v2.5.0).

### Analysis of ALY loci sequence homology

ALY loci were defined based on ALY gene clusters found on the *N*. sing1 genome assembly. 30 ALY gene loci were assigned in total, comprising 91 ALY genes. To analyze these ALY loci, sequences were extracted with 2.5 kb extensions (flanks) on each end or until the ends of the contig for loci situated near contig terminals. Loci sequences were self-aligned using the Dotter software (v4.28)^64^. Homology between ALY genes and intergenic sequences within a locus was identified visually based on this output (Supplementary Fig. 5). Sequence homology for flank, genic and intergenic regions was measured through sequence percent similarity using the Needleman-Wunsch global alignment algorithm with modified parameters (Fig. 3b and Supplementary Fig. 6b). Specifically, pairwise comparisons between these regions were made for each locus using the Biostrings package (v3.18) in R (v4.2.2). Gap opening and extension penalties were increased (gapOpening = 25, gapExtension = 10) to obtain alignments with contiguous regions of high sequence similarity. End gap penalties were omitted to account for differences in lengths (Parameter: type = “overlap”), and the EDNAFULL substitution matrix was used. Percent similarity scores were calculated using the “PID3” formula (100 * (identical positions) / (length shorter sequence)) included in the Biostrings package.

### DNA amplification and gene cloning

Representative full-length alginate lyase genes (from *N*. sing1 and *Vibrio* sp.) and TARP repeat regions (from *N*. sing1) were synthesized as *NdeI*-*BamHI* fragments in pET15b (GenScript). To express and purify individual A- and C-domains from *E. coli*, domain sequences were sub-cloned into a modified pET15b vector containing a C-terminal TEV-6xHIS sequence. To express and purify proteins as N-terminal mCherry or mGFP-fusions, a pET15b vector containing mCherry- or mGFP-TEV- 6xHIS was used. To purify TARP-mCherry fusions, the mCherry gene was subcloned into the synthesized pET15b::6xHIS-*NdeI*-TARP-*BamHI* vector after the *BamHI* site. To assess the trafficking of signal sequences and transmembrane domains (TMDs), the predicted signal sequence from *N*. sing1 ALY49^A3^ (Ref: Sing2_08162) (SS^49^), *Vibrio casei* alginate lyase (GenBank accession: WP_244913617.1) (SS*^V.casei^*^-ALY^), and the human luminal protein Binding Immunoglobulin Protein (BiP) (SS^BiP^) (Genbank accession: NP_005338.1) was appended to the N-terminal end of a sfGFP or mCherry insert by PCR. The C-terminal end of these inserts was also appended with either the ER retention signal KDEL (for SS^BiP^-sfGFP and SS*^V.casei^*^-ALY^-sfGFP) or the predicted transmembrane domain from ALY49^A3^ (for SS^ALY49^). For expression in HeLa cells, constructs were sub-cloned into the pcDNA3.1+ vector between the *NheI* and *XbaI* restriction sites. All constructs were produced through standard molecular biology techniques. PCR amplification was performed using the KAPA HiFi PCR Kit (Roche Diagnostics, 7958838001) and cloning was performed using the In-Fusion® HD Cloning Kit (Takara Bio, 639650). All accessions, primers and vectors used in this study can be found in Supplementary Table 7.

### Protein expression and purification

All recombinant proteins were expressed in BL21 DE3 *E*. *coli* and purified as previously described^135^ with some modifications. Briefly, overnight grown cultures were diluted 1:10 into fresh LB media (supplemented with 100 μg/mL ampicillin) at 37°C, and expression was induced with 1 mM isopropylthiogalactoside (IPTG) once the culture had reached an OD^600^ of 0.6-0.8. Expression was carried out for 16-18 h at 16°C (for ALY48^A5C5^-A1) or 25°C with shaking at 225 rpm. Cultures were then harvested, washed and cell pellets frozen at -80°C. Cell pellets were resuspended in five volumes of lysis buffer and lysed by sonication on ice. Insoluble material was separated by centrifugation and the supernatant fraction was incubated with HisPur Ni-NTA resin (Thermo Scientific, 88222). Bound resin was washed 3 times before elution in lysis buffer containing 500 mM imidazole. Protein fractions were then pooled and dialyzed overnight in 1×TBS (10 mM Tris, 150 mM NaCl) at 4°C using 3,500 MWCO dialysis tubings (Spectrum Laboratories, 25219-041). Following dialysis, proteins were concentrated using 3,000 MWCO centrifugal filters (Merck, UFC500396) and protein concentrations were quantified by Bradford Assay (Bio-Rad, #5000006). For purification of CBM32-mCherry recombinant proteins, cell lysis and elution were performed in a denaturing buffer as previously described^135^. Protein re- folding was performed by dialysis in denaturing buffers containing decreasing levels of urea (4M, 2M and 1M) for an hour each before exchanging into 1×TBS overnight.

### Alginate liquefaction assays

Alginate liquefaction-vortex assays (Fig. 4a) were performed using crude lysates of *E*. *coli* BL21 DE3 harboring ALY pET15b expression plasmids. Alginate gels were prepared by adding 20 mL 2 % (w/v) medium-viscosity sodium alginate (Sigma-Aldrich, A2033) to 20 mL 1×TBS-Ca^2+^ buffer (10 mM Tris, 150 mM NaCl, 2 mM CaCl2) and mixed by shaking. The resulting mixture was briefly centrifuged at 1,000 × *g* for 5 minutes to create a uniform gel. To set up the assay, alginate gel was pipetted into a 15 mL polystyrene tube (Falcon®, Corning) until the 2 mL mark and 80 μL cell lysate was added onto the gel surface. Reactions were incubated at 30°C overnight. To quantify gel liquefaction, tubes were vortexed from the bottom for 5 s at 1.5 speed setting on touch using a vortex mixer (Vortex-Genie 2 SI-0236, Scientific Industries Inc) and the height of liquid displacement was marked and measured manually. All assays were performed in triplicates.

### Alginate lyase enzyme activity assays

Alginate lyase activity was determined spectrophotometrically at 235 nm by following the formation of 4,5-unsaturated bonds at the non-reducing end^136,137^ (Fig. 4b). Purified recombinant A-domains were added at a final concentration of 0.1 μM to an alginate lyase enzyme activity buffer (0.2 % (w/v) low-viscosity sodium alginate, 100 mM Tris, 2 mM CaCl2) at a reaction volume of 200 μL. Purified mCherry-6xHIS and a commercial alginate lyase (Sigma-Aldrich, A1603) serve as negative and positive controls, respectively. 1×TBS was used as blank. Assays were prepared in triplicates in a 96-well UV-transparent microplate (Corning®) and measurements were taken at 25°C for 5 minutes (every 5 s interval) using a Tecan Spark Multimode Microplate Reader (Tecan Inc).

### Transient transfection of signal sequences

To assess the trafficking of signal sequences, vectors containing the signal sequence for *V*. *casei* or a positive control consisting of the signal sequence from the human luminal protein BiP were transfected into HeLa mammalian cells and imaged as previously described^138^ (Fig. 1d). HeLa cells were transfected with plasmids using the lipofectamine™ 3000 transfection reagent (Invitrogen, L3000001) and cultured for 48 h. Imaging was performed on the SP8 Inverted gSTED confocal microscope (Leica Microsystems) fitted with a 100× objective (NA: 1.4). To monitor the trafficking signals derived from ALY49^A3^, the SS^49^-mCherry-TMD^49^ construct was co-transfected with the ER marker SS^BiP^-sfGFP-KDEL. Images were taken at 24 and 72 h post-transfection. For imaging, cells were cultured and transfected inside a 35 mm petri dish with a 20 mm glass bottom (VMR, 734-2906). Lysotracker-DND-26 (Invitrogen, L7526) was used to visualize lysosomes in transfected HeLa cells at a final concentration of 50 nM (Fig. 5f). Lysotracker was added directly to the medium in the dish and incubated for 15 minutes at 37°C. The probe-containing medium was then removed and replaced with 1×PBS prior to imaging.

### Vacuole staining and fluorescence measurement

Fluorescent alginate moieties that appear upon growth of *N.* sing1 (Fig. 5a and c) were visualized using epifluorescence and a CFP filter set (excitation 434 nm, emission 479 nm). Diatom vacuoles were stained using the CellTracker™ Blue CMAC dye (7-amino-4-chloromethylcoumarin) (Invitrogen, C2110) at a working concentration of 100 μM (Fig. 5c and Supplementary Fig. 11a). For liquid cultures, CMAC was added directly into the medium and incubated at room temperature for 15 minutes. Cells were washed with an equal volume of SSW medium three times before imaging. For solid cultures, 10 μL 100 μM CMAC was added onto a 1×1 cm square agar block of diatom culture and incubated at room temperature for 15 minutes. To obtain and compare the emission spectra between vacuoles from alginate-grown diatoms (3 d) and the culture supernatant (4 d), samples were excited at 405 nm and scanned between 410-710 nm (step width: 20 nm). Intensities were normalized between 0-1. Data shown is derived from three independent scans (Fig. 5e). Absorbance wavescans of diatom and *Vibrio* sp. culture media were performed using a Tecan Spark Multimode Microplate Reader (Tecan Inc) (Fig. 5b).

### Alginate substrates

The alginates purchased from Sigma-Aldrich (A2033) and Pronova (42000101; 42000301) were analyzed using the standard protocol outlined by ASTM (American Society for Testing and Materials) method F2259-10 (2012) to determine their chemical composition, including sequential parameters, via NMR spectroscopy. Additionally, the molecular weight of the alginates was determined using SEC-MALS following the standard protocol provided by ASTM method F2605-08. For both methods, the analytical protocol for alginate^139^ has been previously described. Poly-M was obtained from an epimerase-negative AlgG mutant of *Pseudomonas fluorescens*^139^. poly-G and poly-MG were prepared as previously described^140^. Hydrolyzed seaweed alginate was prepared using an alginate sample extracted from *Laminaria hyperborea* leaf and subjected to stepwise acid hydrolysis to achieve an approximate degree of polymerization (DP) of 30. Initially, 1 % (w/v) alginate was dissolved in ion-free water, with the pH adjusted to 5.6 using 0.1 M HCl, followed by hydrolysis at 95°C for 1 hour. Subsequently, the solution was cooled to room temperature in a water bath, and the pH was adjusted to 3.8. The hydrolysis was then continued at 95°C for 50 minutes. After cooling again to room temperature in a water bath, the solution was neutralized to pH 6.8–7.5 using 0.1 M NaOH and then lyophilized.

### Alginate-binding assays

Alginate microgels were prepared by mixing 1 % (w/v) medium-viscosity sodium alginate with 10 mM CaCl2. The mixture was then centrifuged at 3,000× *g*, 4°C, for 15 minutes and resuspended in 1×TBS-Ca^2+^. To perform the binding assay, 3–5 μM of the mCherry-fusion protein was incubated with 1 mg/mL bovine serum albumin (BSA) (Sigma-Aldrich, A3059) in 1×TBS-Ca^2+^ for 15 minutes at room temperature (Supplementary Fig. 14a). For binding assays involving both TARP-mCherry and CBM32-mGFP (Fig. 6b and Supplementary Fig. 13b), proteins were incubated at a 1:3 ratio with 2 mg/mL BSA in 1×TBS-Ca^2+^ for 15 minutes at room temperature. Following the incubation, proteins were centrifuged at 21,130× *g* for 10 minutes. The supernatant fraction was then mixed with the alginate microgel in a 1:1 ratio, vortexed and incubated at room temperature for another 30 minutes. Afterwards, this microgel- protein mixture was centrifuged at 21,130× *g* for 5 more minutes. The resulting alginate protein pellet was then washed twice with 1×TBS-Ca^2+^. 5 μL of the mixture was used for fluorescence microscopy (Olympus BX51, Olympus). For alginate-binding assays involving soluble alginates (Fig. 6c), proteins were added (1:10 dilution) with an alginate solution (0.3% medium-viscosity sodium alginate, 1 mg/mL BSA, 1×TBS), vortexed and diluted 64× before imaging under fluorescence microscopy.

FRAP (Fluorescence recovery after photobleaching) was used to assess the binding of CBM32 domains from *N*. sing1 ALY77 and *Vibrio hyugaensis* (Supplementary Fig. 14b). Alginate binding was performed using the alginate microgels as described above, and samples were imaged using an Olympus FV3000 Inverted confocal system (Olympus) at 40× magnification N.A. of 1.4. Four frames were taken before samples were bleached with 100 % laser power at maximum speed for one frame (fifth frame) before recovery (frames taken every 2 s for up to 300 frames). FRAP analysis was carried out using the cell-Sens software (Olympus). Normalization was carried out against frames 1-4.

Alginate binding was also assessed with FPLC (ÄKTA Pure™ FPLC system, Cytiva) (Fig. 6c). 4 μM TARP-mCherry recombinant protein was mixed with 0.1 % low-viscosity sodium alginate and incubated in 1×TBS+ (10mM Tris, 200 mM NaCl) for 15 minutes prior to sample injection. mCherry-6xHIS was used as a negative control. Mixtures were separated in 1×TBS+ through a Superdex 200 Increase 10/300 GL (GE Healthcare, 28990944) column. Protein peaks were monitored at 260 nm and 587 nm.

### Sargassum staining

*Sargassum* material was collected at low tide from the intertidal zone on Sentosa island, Singapore (latitude 1.259895, longitude 103.810843; Singapore National Parks Board permit no. NP/RP20-016). *Sargassum* fronds were washed in SSW medium and stored in 70 % ethanol to extract and remove the chlorophyll. Once coloration was sufficiently lost, 1×1 cm squares were cut from the edge of the frond along the sagittal plane to produce thin strips of tissues. To minimize non-specific binding, strips were blocked with an SSW-based buffer containing 10 mg/mL BSA and 0.1 % Tween 20 for 30 minutes. After blocking, tissues were washed with SSW and incubated with the recombinant proteins for 1 h. Following this, strips were washed in 200 μL SSW three times for 10 minutes each. The cross-section of the tissue was imaged using fluorescence microscopy (Olympus BX51, Olympus) at 100× magnification (Supplementary Fig. 14c).

### NMR analysis and time-resolved NMR

NMR spectra were acquired at 25°C on a BRUKER AVIIIHD 800 MHz or Bruker AVIIIHD 600 MHz spectrometer (Bruker BioSpin AG, Fällanden, Switzerland) both equipped with a 5 mm z-gradient CP-TCI (H/C/N) cryogenic probe. The chemical shift was referenced to residual water signal ^1^H: 4.75 ppm and the ^13^C chemical shift was referenced indirectly to water, based on the absolute frequency ratios^141^. Spectra were recorded, processed, and analyzed using TopSpin 3.5 or 4.1.4 (Bruker BioSpin). Reactions were run in 200 μL 20 mM HEPES pH 7.5, 25 mM NaCl, 2 mM CaCl2 in D2O (D, 99.9 %) in 3 mm LabScape Stream NMR tubes (Bruker LabScape).

For time-resolved experiments, 10 mg/mL seaweed alginate, poly-M, poly-MG or poly-G substrate (Supplementary Table 8) was dissolved in the HEPES buffer. 1D proton spectrum with water suppression (noesygppr1d) was recorded at 25°C to verify the sample’s integrity before the time-resolved NMR experiment. The reaction was started by adding 1-2 μL of ALY7-A (50 μM), ALY48^A5C5^-A1 (60 μM), ALY58-A (50 μM) or ALY77-A (50 μM) to the preheated substrate and mixed by inverting the sample a few times. The sample was then immediately re-inserted into the NMR instrument and the experiment started. The recorded spectrum is a pseudo-2D type experiment recording a 1D proton NMR spectrum (based on noesygppr1d) every 5 min with a total of 64 time points (total experiment time 5 h 20 min). After each time-resolved experiment, a ^1^H-^13^C HSQC (heteronuclear single quantum coherence; hsqcetgpsisp2.2) spectrum with multiplicity editing was recorded. Signals were assigned based on previously published assignments from^132,142–144^ (Fig. 4d-e and Supplementary Fig. 10).

## Acknowledgements

Work in the Jedd Group is supported by the Temasek Life Sciences Laboratory. FLA acknowledges the Norwegian Research Council for funding the NNR (Norwegian NMR Platform) (226244), SBP-N (294946), and AlgModE (315385). MN acknowledges funding by the German Research Foundation (DFG, NO407/7-2). FLA is grateful for experimental assistance from Synnøve Strand Jacobsen and Alexander Mika Hannasvik. We thank Mirjam Czjzek and Antonia Monteiro for helpful discussions.

## Author Contributions

G.J. conceived the project. M.N. performed genome assembly and annotation. Z.H.L. performed phylogenetic and other bioinformatics analyses. F.L.A. performed NMR analyses. Z.H.L., P.Z., C.Q. and G.J. performed the experiments. G.J. and Z.H.L. wrote the manuscript with input from all authors.

---

**Supplementary Figure 1.**
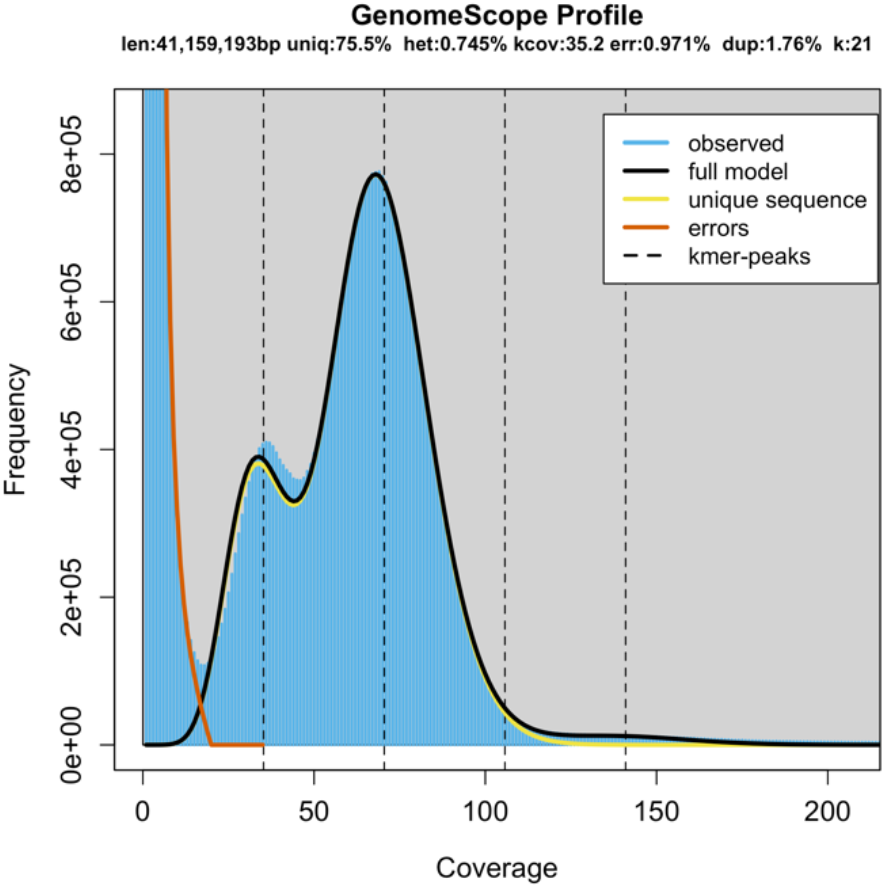
N. sing1 is likely to be diploid. 21-mers (*k*-mers) were counted using the tool Jellyfish (v2.3.1). The presence of two major peaks observed in the GenomeScope profile suggests that the *N*. sing1 genome is diploid.

**Supplementary Figure 2.**
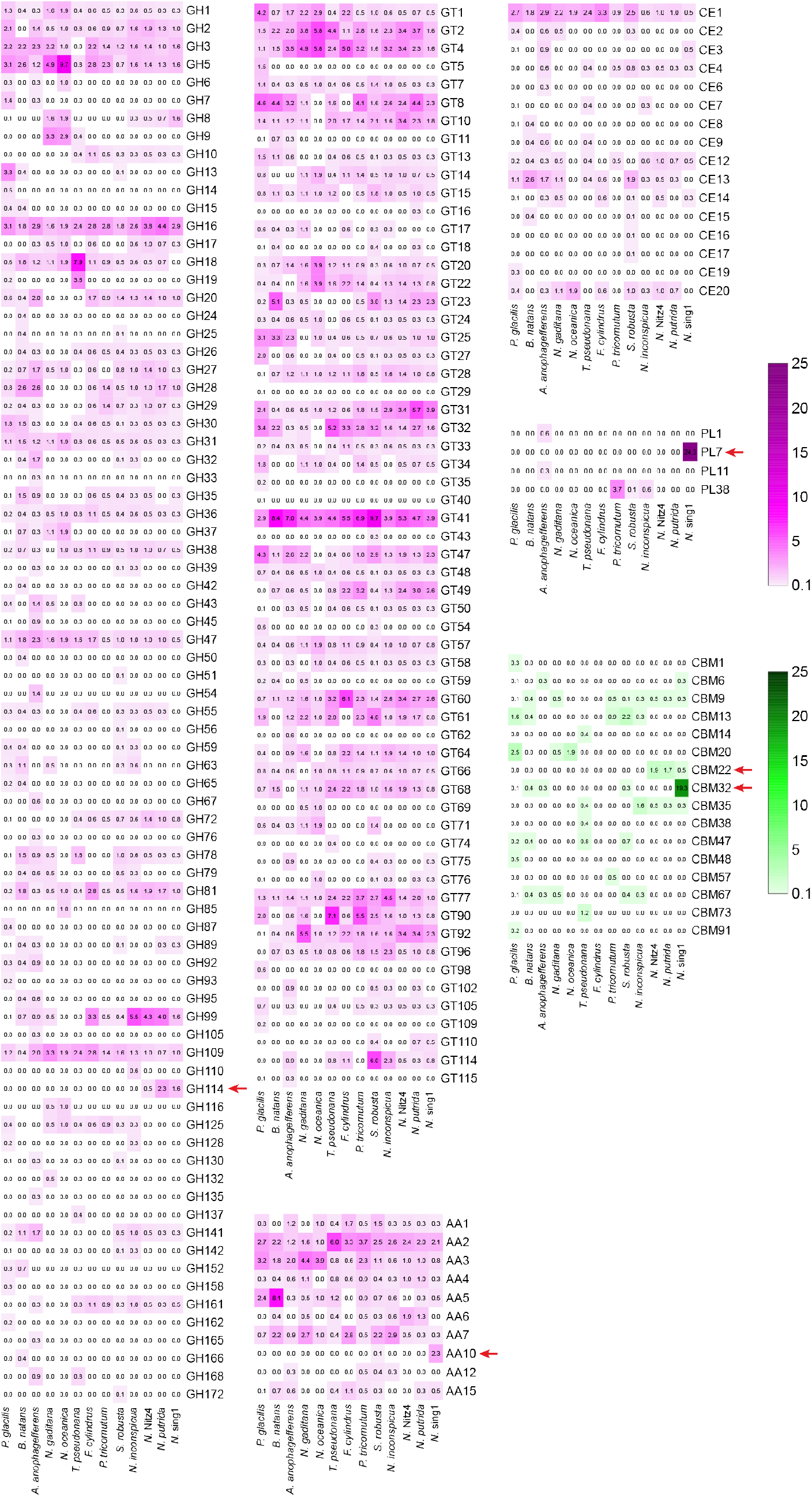
Complete carbohydrate-active enzyme (CAZyme) analysis. The heatmap displays the prevalence of different CAZyme (enzyme classes) families (magenta) and carbohydrate-binding domain (CBM) families (green) annotated in the protein-coding genes of each taxon. Values for each family are calculated as a proportion of the total CAZyme count (enzyme class only) detected in the protein-coding genes of each taxon and are reflected through color intensity. Red arrows denote families that appear to have undergone expansion in the apochlorotic or *N.* sing1 lineage.

**Supplementary Figure 3.**
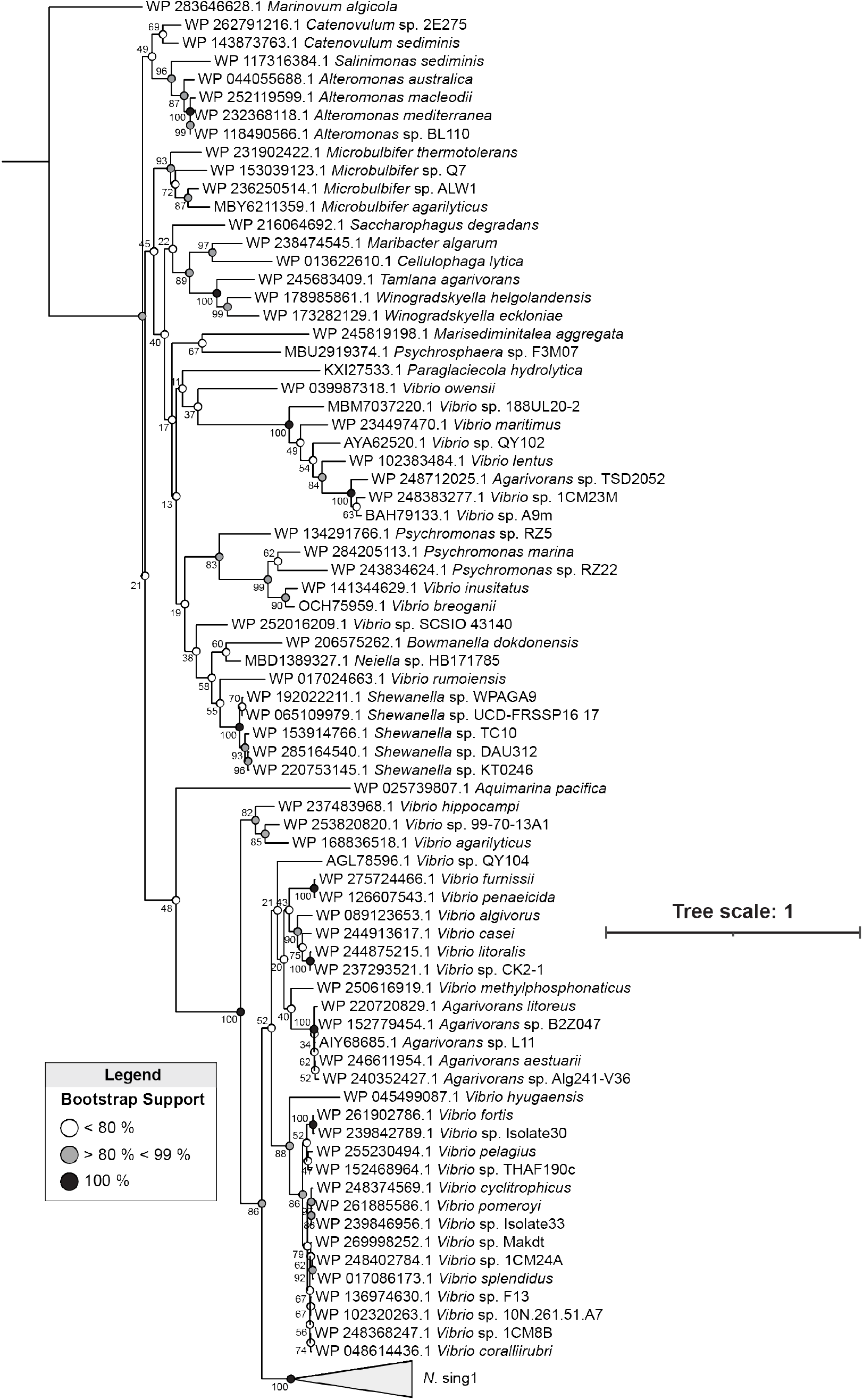
Maximum likelihood phylogeny of alginate lyase domains from bacteria and *N*. sing1 ALY genes. The phylogenetic tree was constructed using a protein sequence alignment of all the alginate lyase domains (A-domains) (bootstrap replicates, n = 1000) from *N.* sing1 and selected bacteria alginate lyases. Branch node support values are indicated and summarized as described in the figure legend. Tree scale = 1. The 91 *N*. sing1 ALYs form a monophyletic clade and are collapsed for illustration.

**Supplementary Figure 4.**
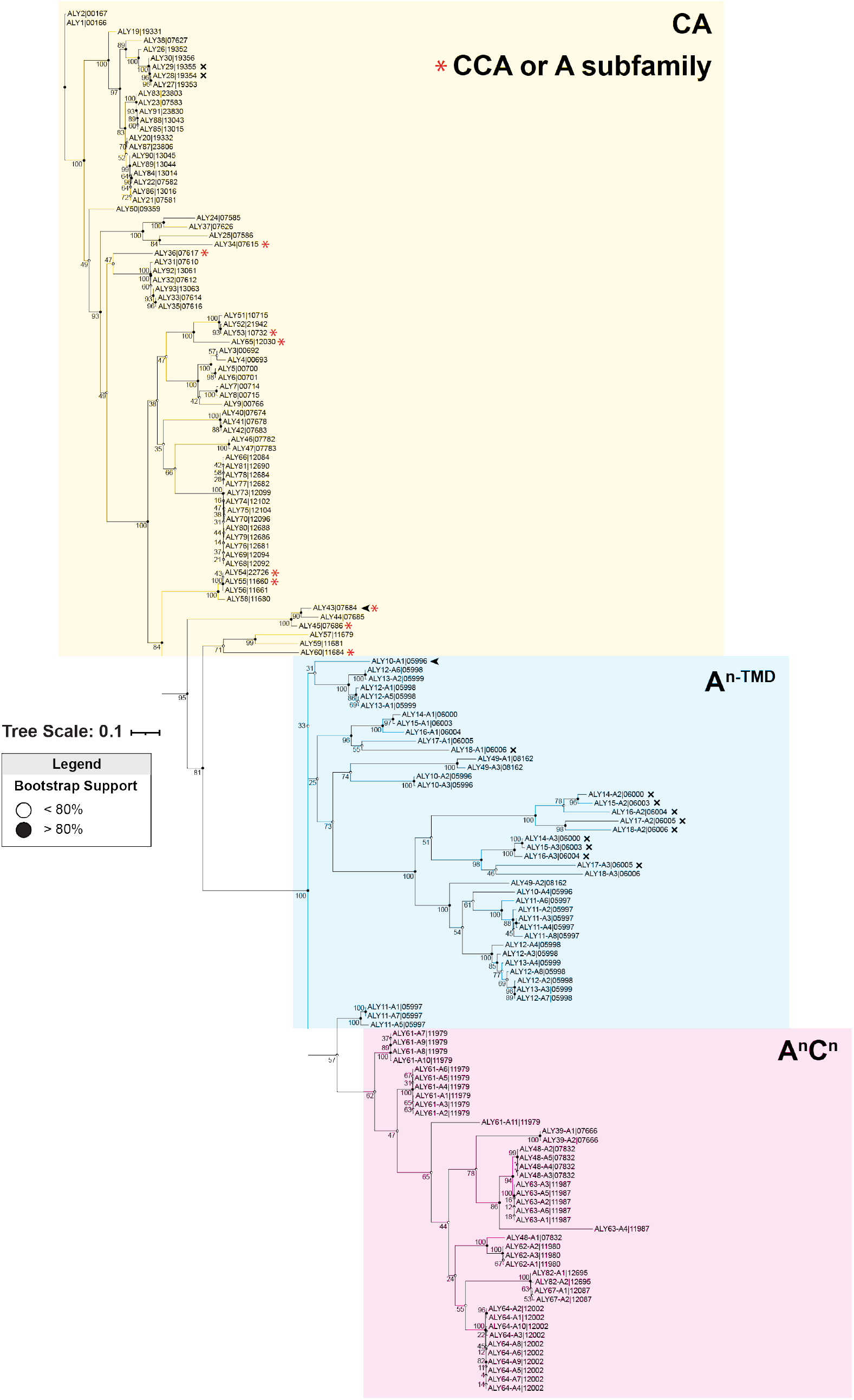
Maximum likelihood phylogeny of alginate lyase domains from *N*. sing1 ALY genes. The phylogenetic tree was constructed using a nucleotide sequence alignment of alginate lyase domains (A-domains) (bootstrap replicates, n = 1000) from *N.* sing1 ALYs. Branch node support values are indicated and summarized as described in the figure legend. Tree scale = 0.1. Clades highlighted in yellow, blue and red fall under the CA, A^n-TMD^, and A^n^C^n^ families, respectively. Red asterisks denote CA family genes with diverged domain organizations (CCA or A subfamily).

**Supplementary Figure 5.**
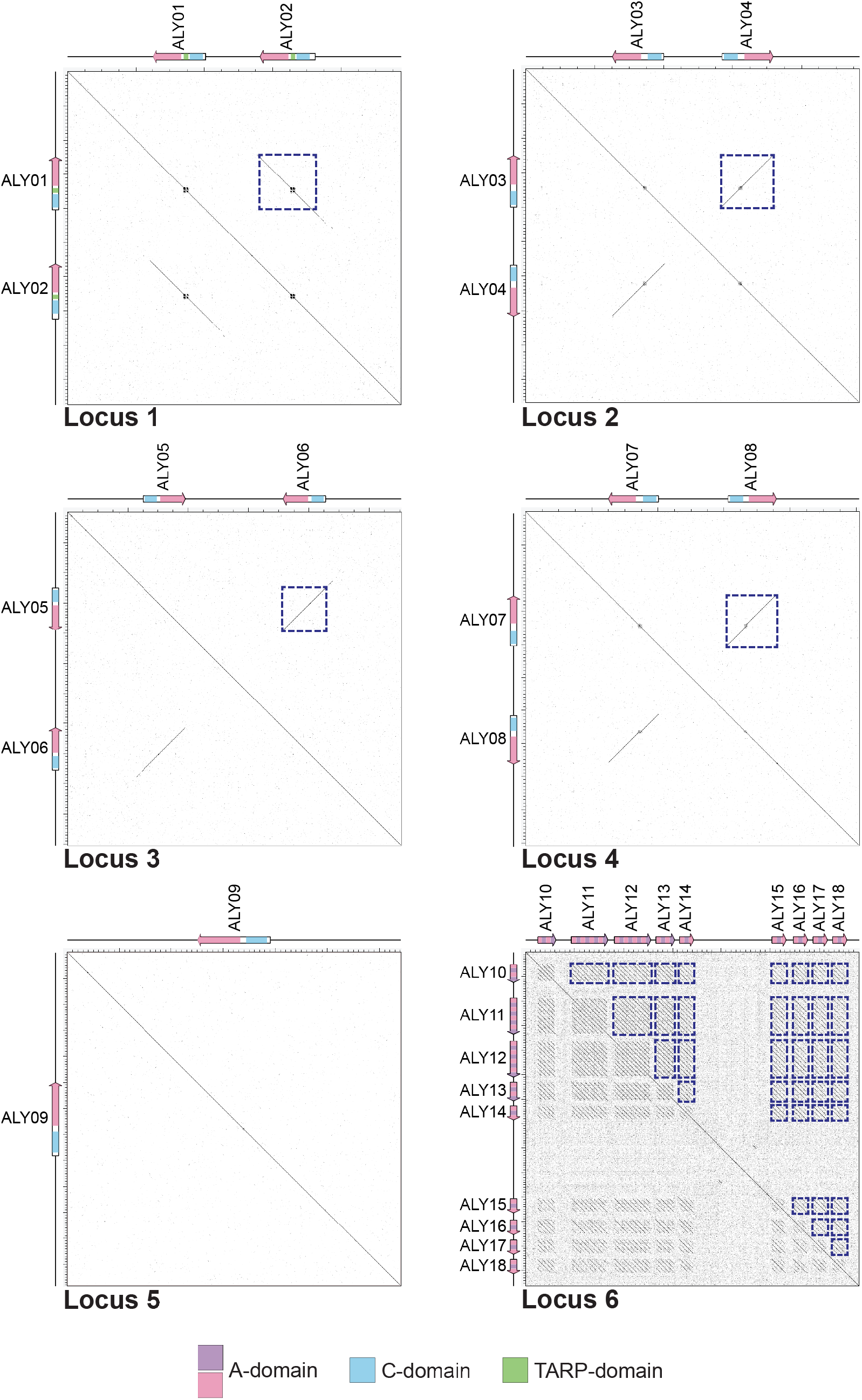
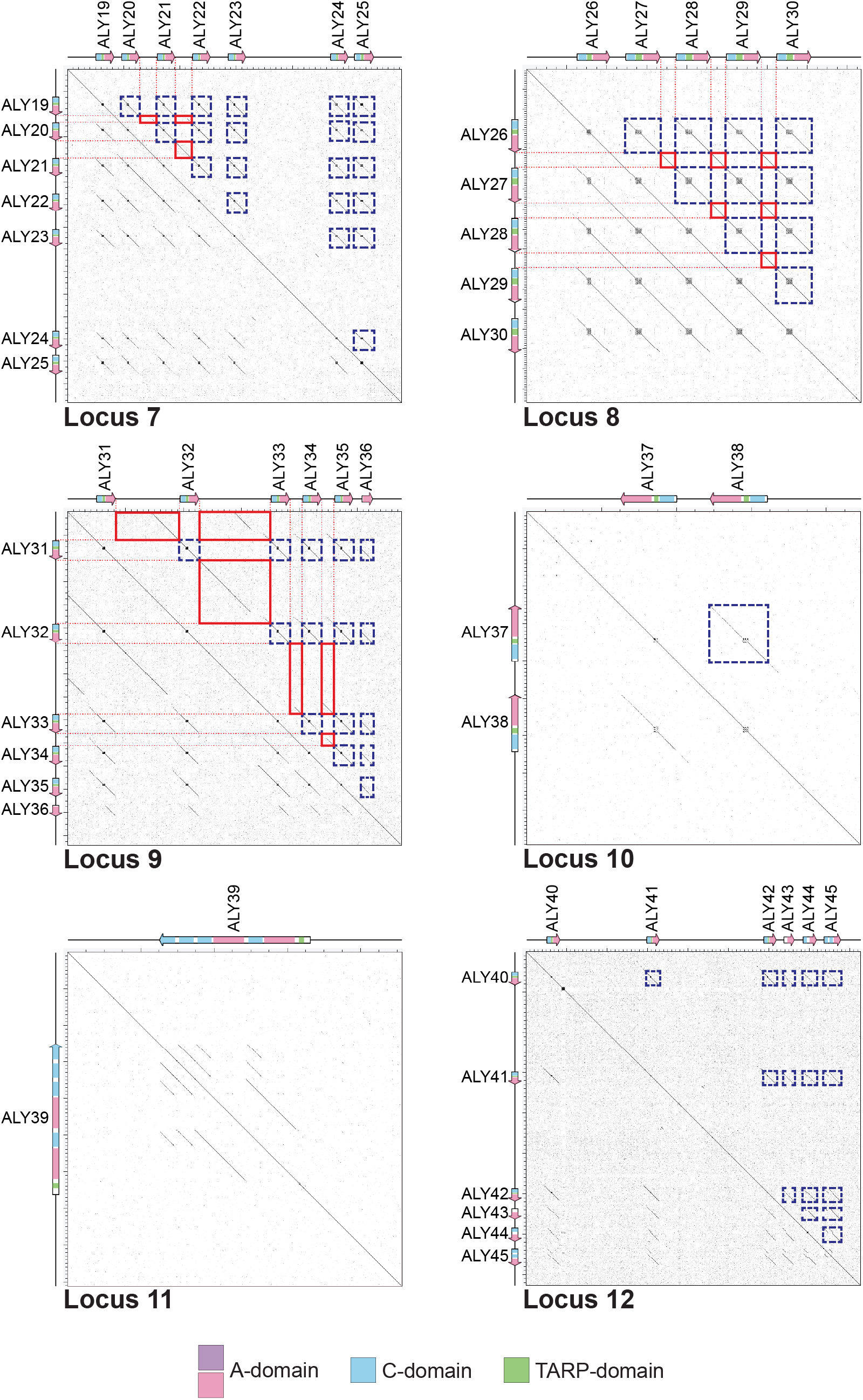
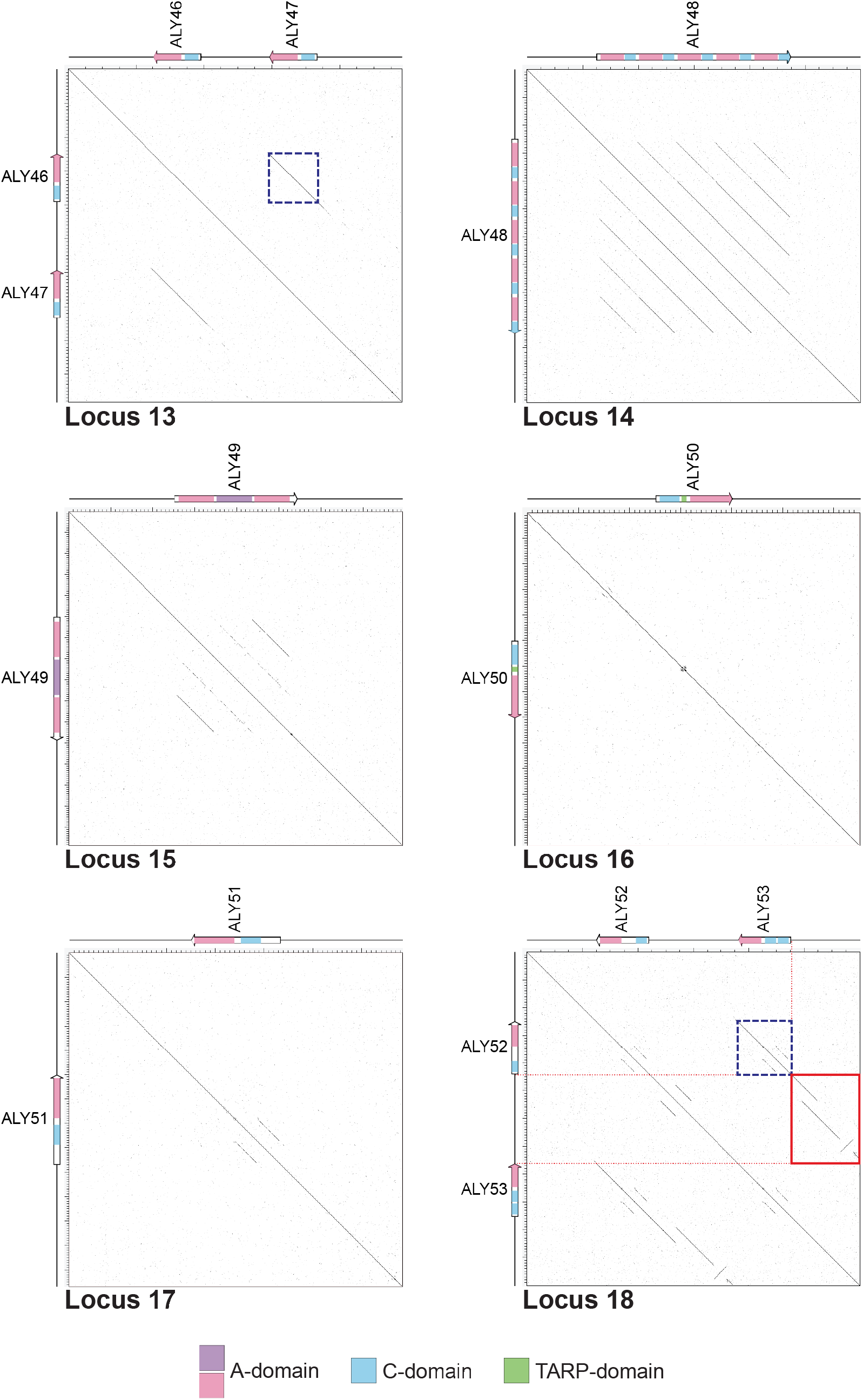
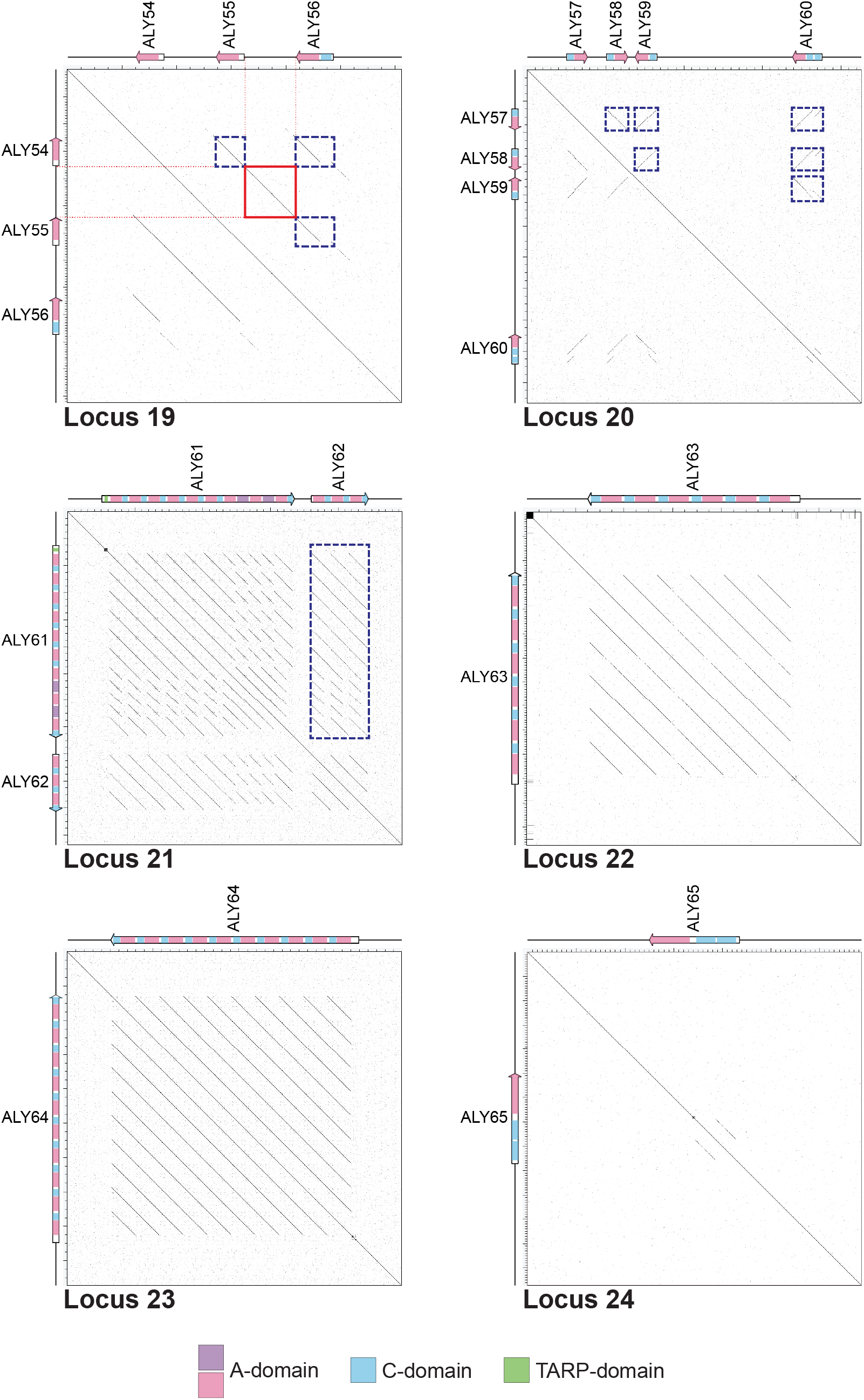
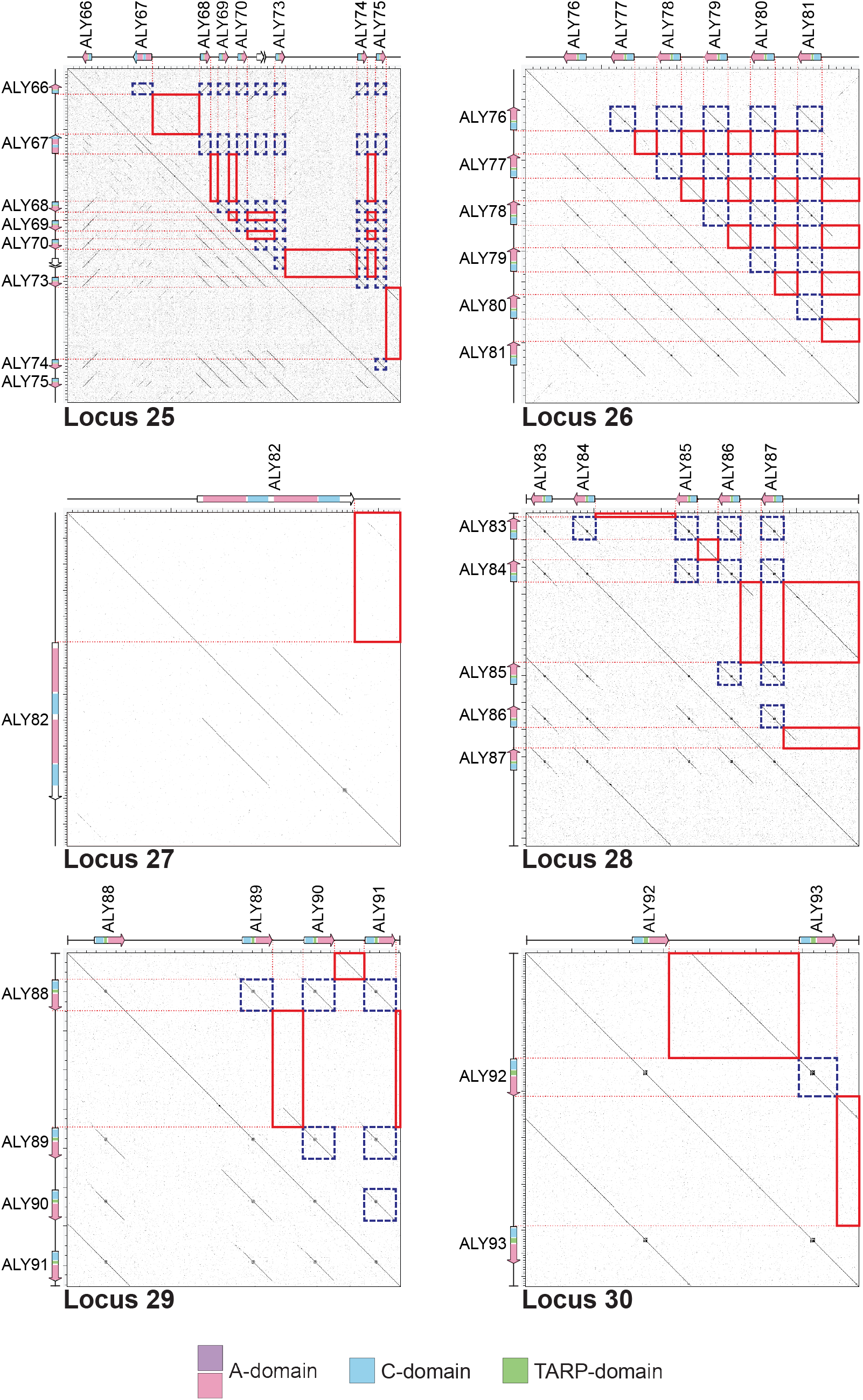
Sequence dot plots of ALY loci in the *N*. sing1 genome. Self-alignment of ALY loci nucleotide sequence (± 2.5 kb) reveals patterns of DNA homology within the loci. The gene structure of the ALY locus is shown along both axes and protein domains are labelled according to the legend. Dashed dark blue and red boxes identify homology between genic and intergenic regions along the locus, respectively.

**Supplementary Figure 6.**
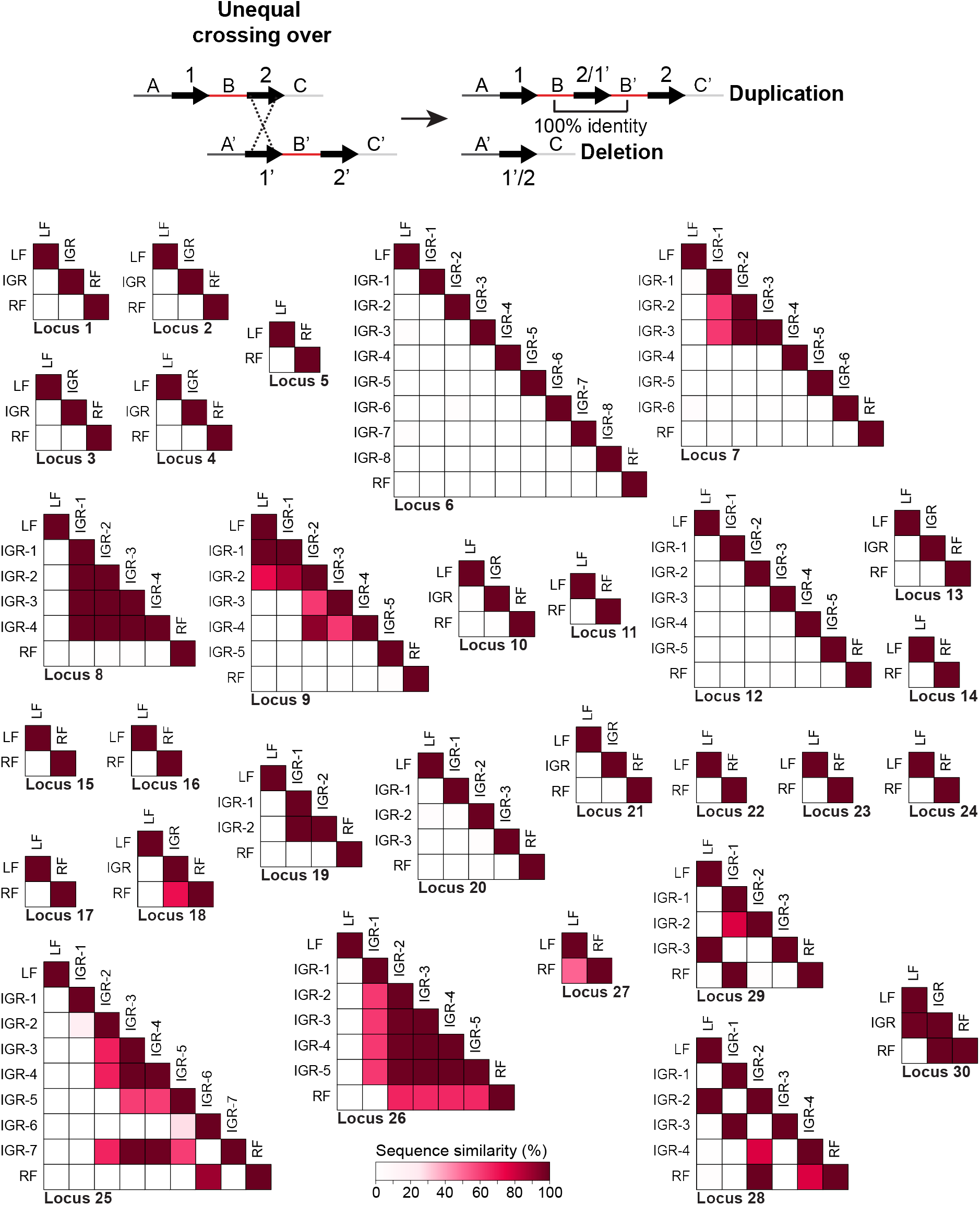
Pairwise comparisons of intergenic sequences in ALY loci of *N*. sing1. (Top) Diagram of an unequal crossing-over event, illustrating how intergenic regions can be duplicated as a result of tandem duplications. (Bottom) Pairwise comparisons between intergenic regions within ALY loci. The sequence percent identity is calculated against the shorter sequence using the Needleman-Wunsch alignment algorithm with modified parameters. Similarity scores are represented as diagonal matrices and reflected by color intensity. IGR: Intergenic Region; LF: Left Flank; RF: Right Flank.

**Supplementary Figure 7.**
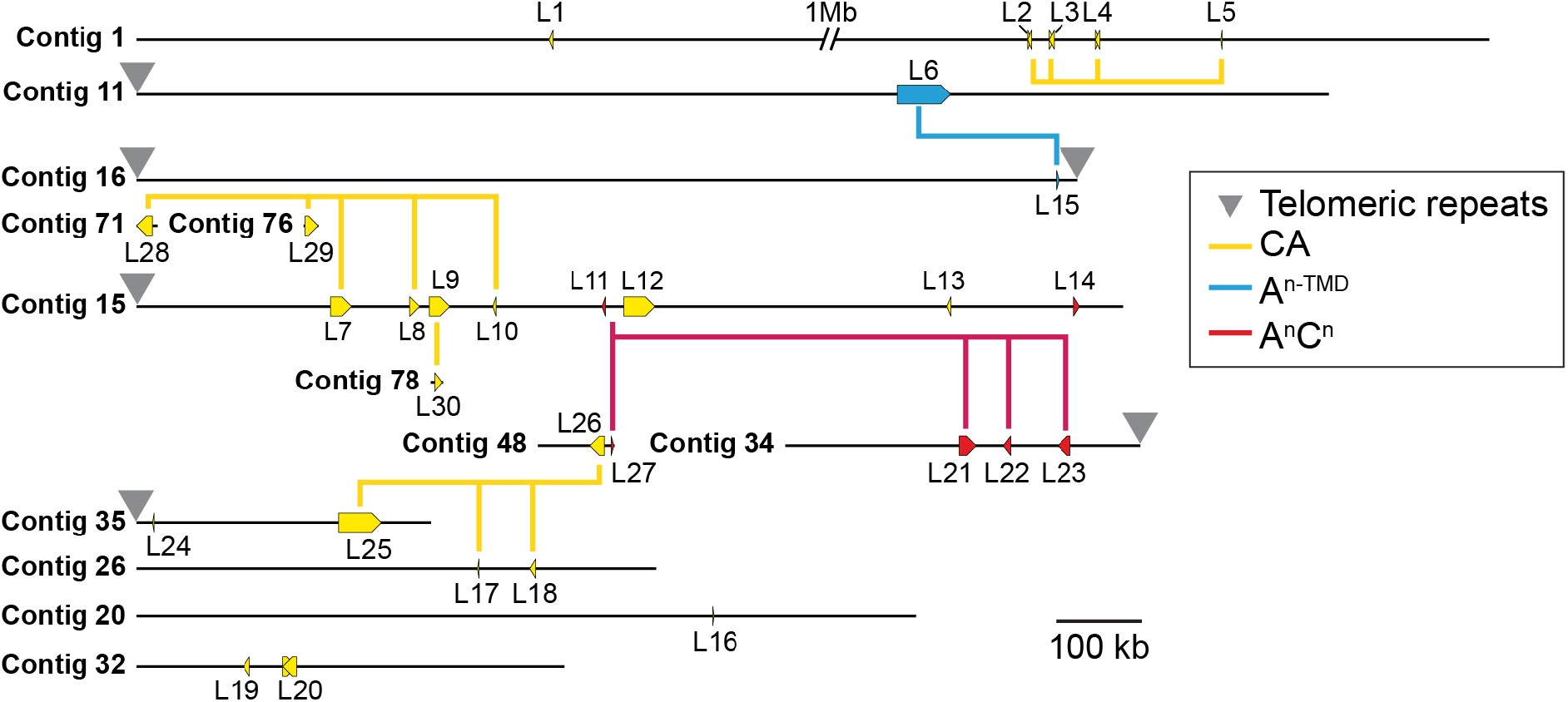
Genomic arrangement of ALY loci in *N*. sing1. Location of ALY loci on contigs of the *N*. sing1 assembly. ALY loci are differentiated by color according to their assigned gene family (yellow: CA; blue: A^n-TMD^; red: A^n^C^n^) and telomeric repeats are identified (gray inverted triangles). Phylogenetically related loci (bootstrap > 99 %) are connected by a line.

**Supplementary Figure 8.**
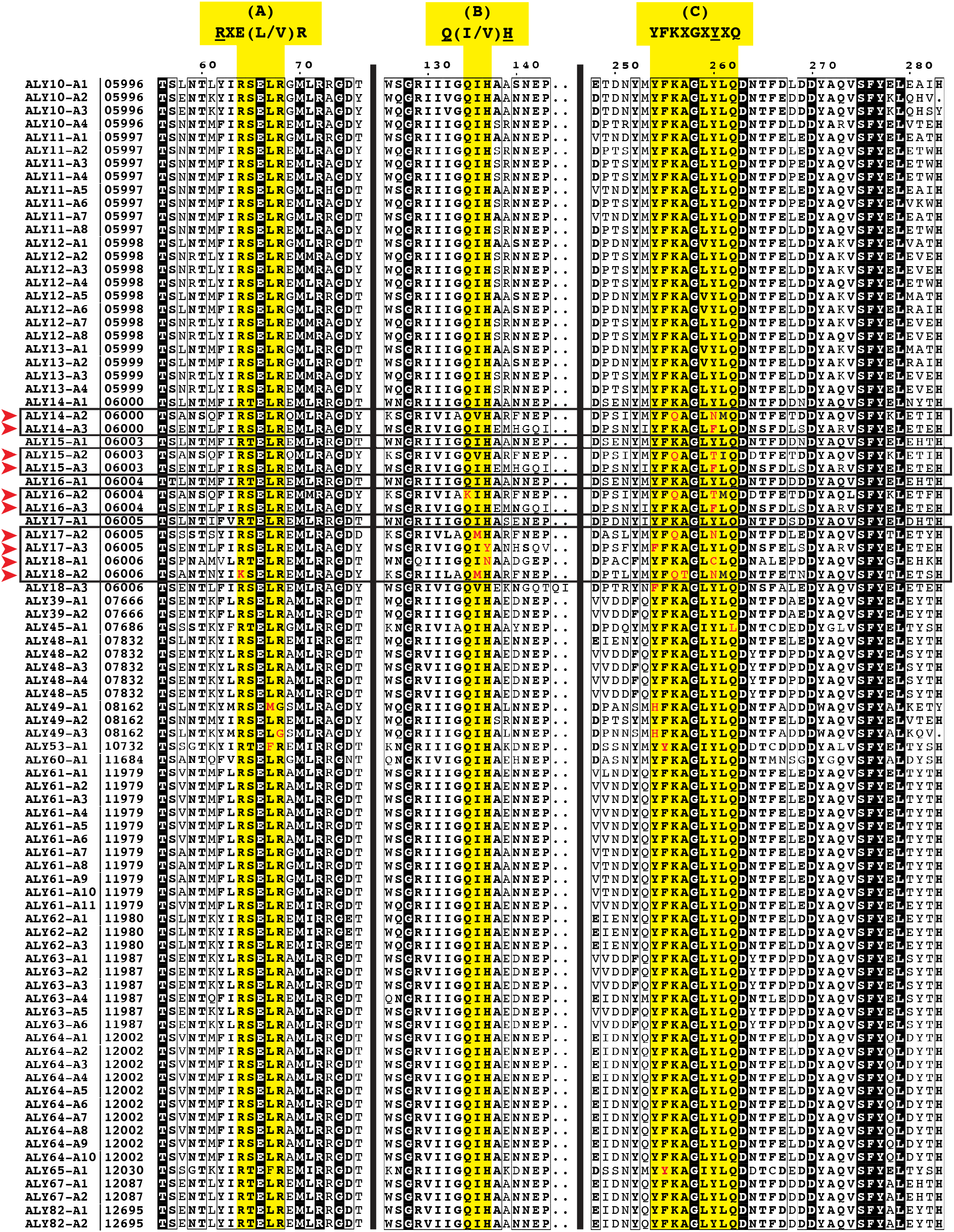
Protein sequence alignment of PL7 catalytic regions in A^n-TMD^ and A^n^C^n^ family ALY proteins. Amino acid sequence alignment of A^n-TMD^ and A^n^C^n^ family proteins in *N*. sing1. Conserved catalytic regions are highlighted in yellow (A: RXE(L/V)R; B: Q(I/V)H; and C: YFKXGXYXQ). The red arrowheads denote proteins with notable substitutions at key catalytic residues in at least one of these three catalytic regions.

**Supplementary Figure 9.**
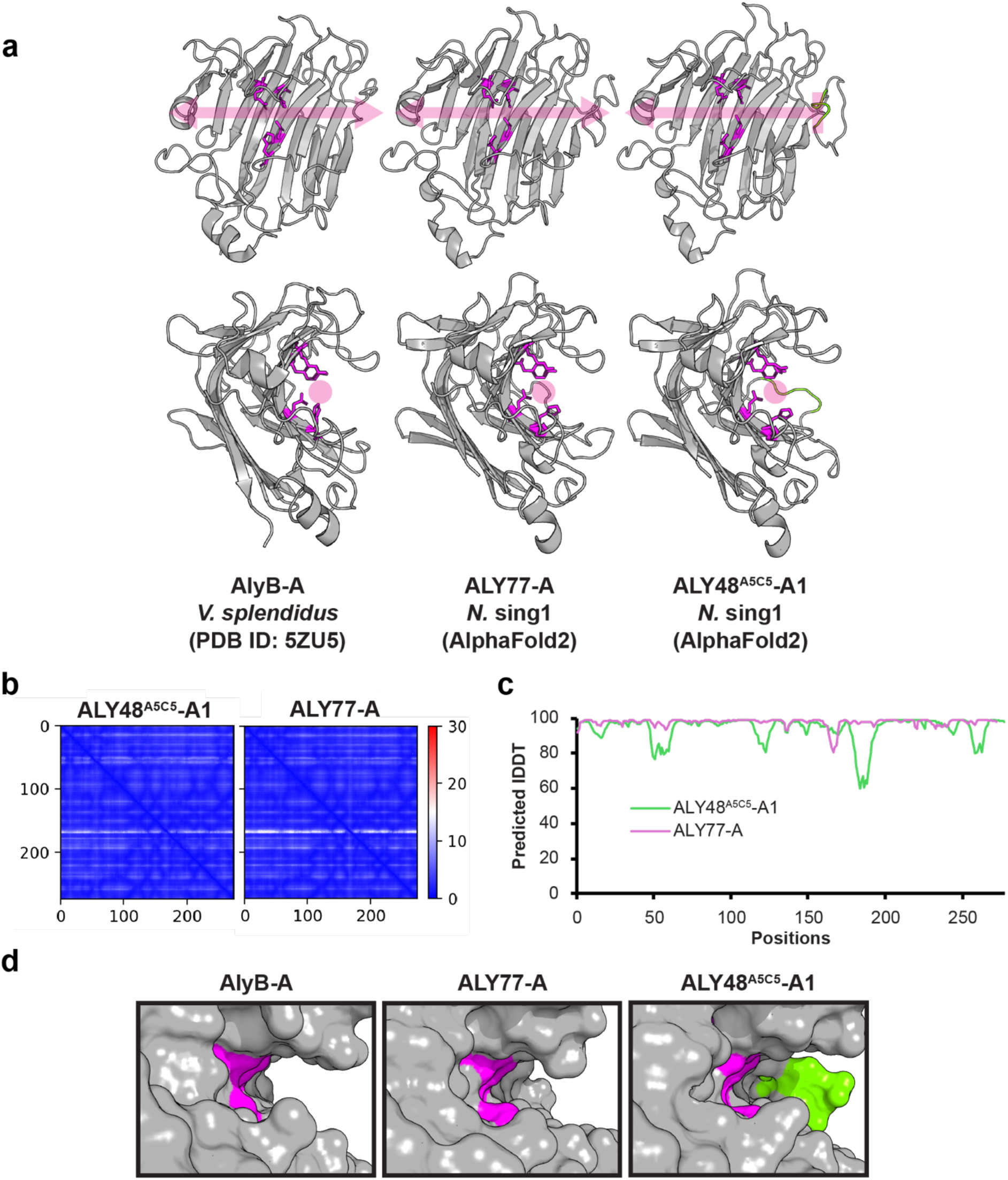
**Structural predictions of *N*. sing1 ALY proteins. a**. (left to right) Cartoon representations of a protein crystal structure from AlyB alginate lyase domain (*V. splendidus*; PDB 5ZU5) and AlphaFold2 predictions of ALY77-A and ALY48^A5C5^-A1 alginate lyase domains (*N*. sing1). The top and bottom rows show two different views along the catalytic cleft. Putative catalytic residues (R/Q/H/Y) are colored in magenta, while insertions predicted to contribute to exolytic activity are colored in green. Gray arrows and circles indicate the likely binding site of alginate molecules along the catalytic cleft. **b**. Predicted aligned error (PAE) plots of the two AlphaFold2-predicted structures in **a** show the confidence in the relative position of two residues within the predicted structure. **c**. Predicted local distance difference test (pLDDT) scores for each residue of the two AlphaFold2-predicted structures in **a**, which signifies the confidence scores of the prediction. **d**. Magnified view of the alginate lyase catalytic cleft displayed as surface representations. Note that the loop insertion in ALY48^A5C5^-A1 appears to form a hindrance on one end of the catalytic cleft.

**Supplementary Figure 10.**
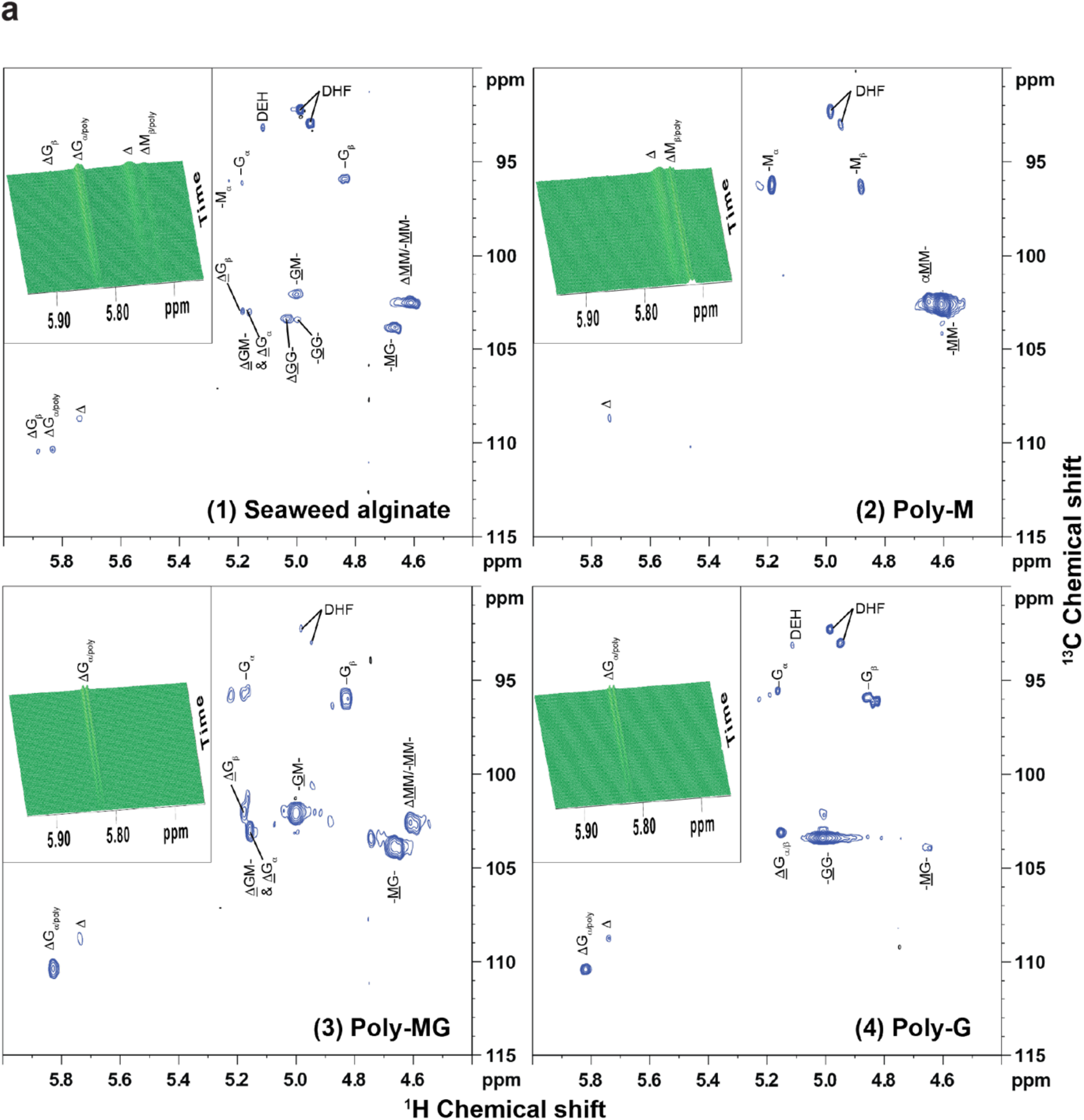
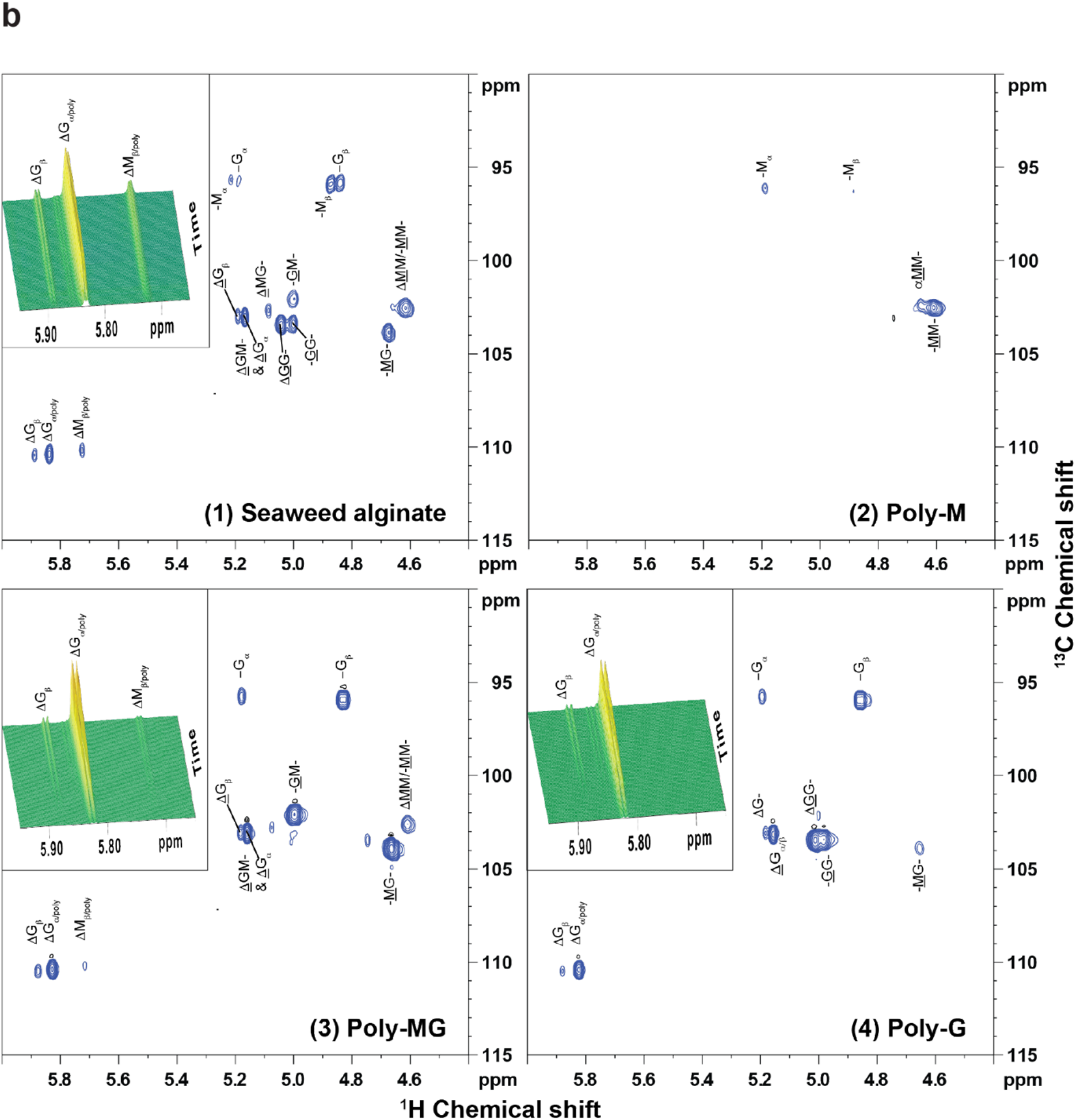
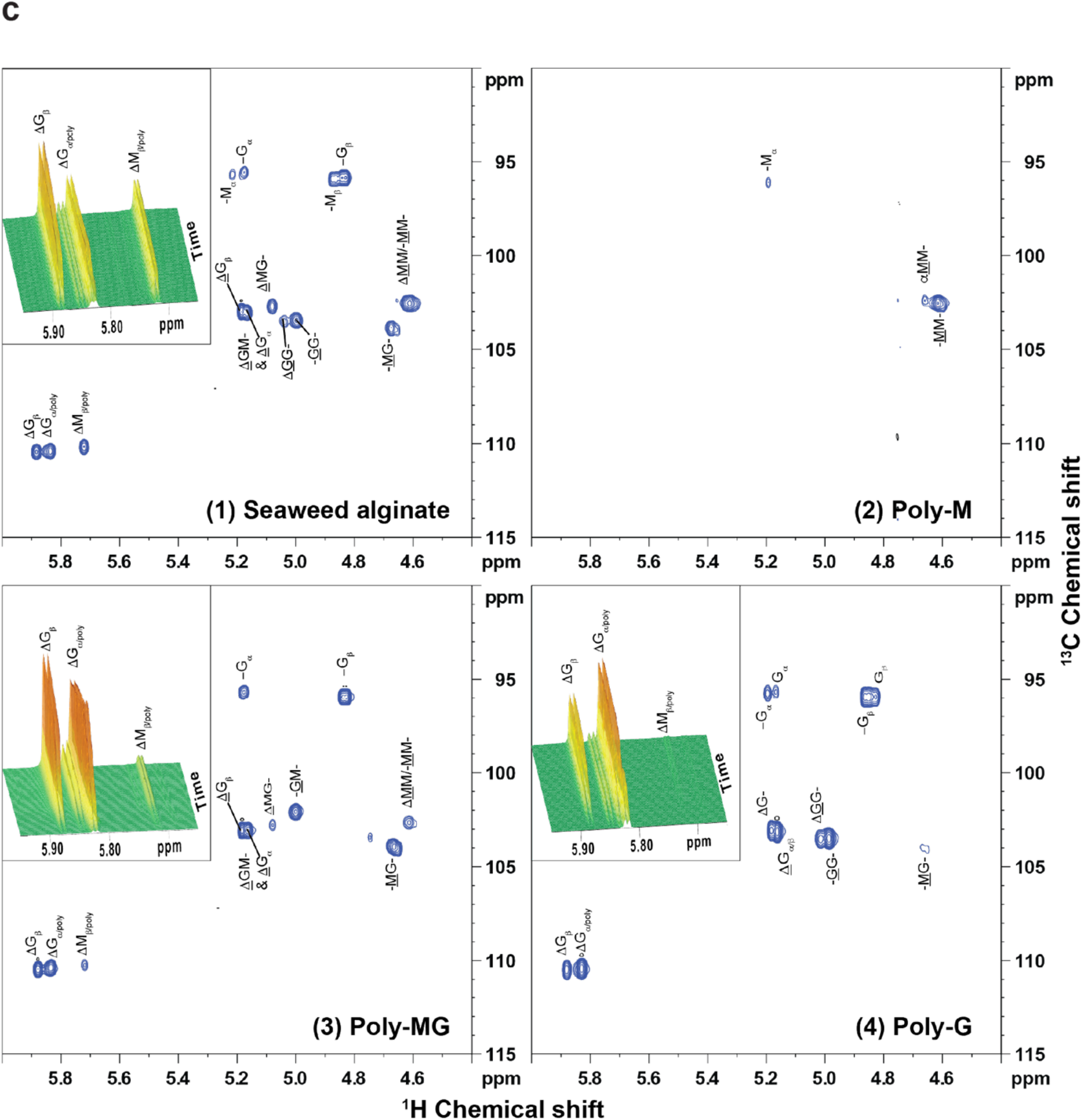
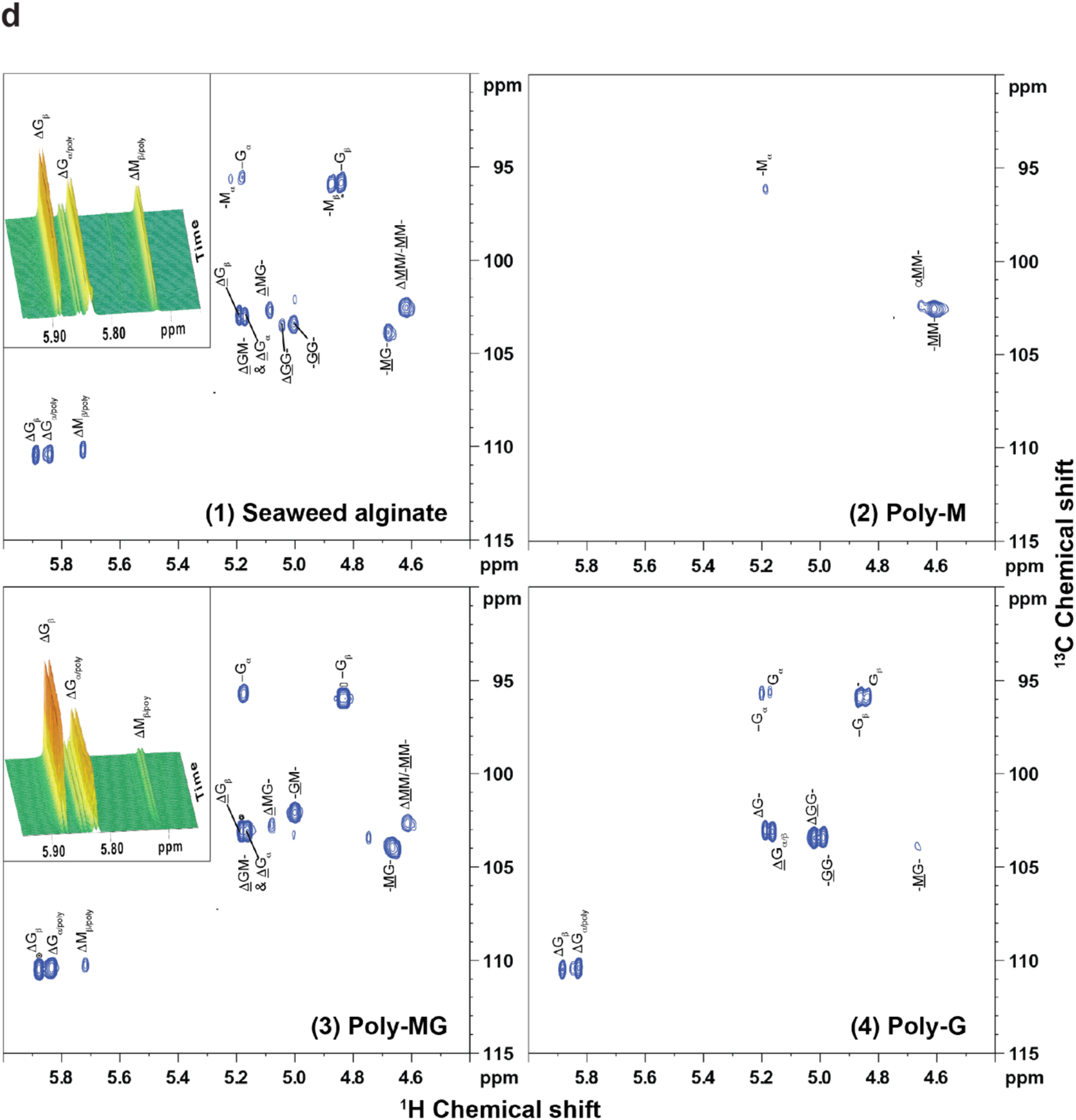
Activities of recombinant *N.* sing1 ALYs on different alginate substrates. Anomeric and ΔH/C-4 regions of ^1^H-^13^C HSQC of (1) Seaweed alginate (FG 0.46; DP ∼30) (top left panel), (2) Poly- M (DP ∼30) (top right panel), (3) Poly-MG (DP ∼26) (bottom left panel), and (4) oligoG (DP ∼26) (bottom right panel) treated with **a.** ALY48^A5C5^-A1, **b.** ALY7-A, **c.** ALY58-A, and **d.** ALY77-A after 5 h 20 min (in 20 mM HEPES (pH 7.5), 25 mM NaCl, 2 mM CaCl2 in D2O (D, 99.9 %) recorded on a 600 or 800 MHz instrument at 25°C. The inlay panels show ^1^H time-resolved spectrum of the ΔH/C-4 region over the reaction period. Inlay panels are omitted in the absence of activity (ALY7-A (2), ALY58-A (2), and ALY77-A (2)) or in cases of errors (ALY77-A (4)—lock error during recording). M: Mannuronate; G: Guluronate; Δ: ΔH/C-4 of 4,5-unsaturated 4-deoxy-L-erythro-hex-4- enepyranosyluronate; <u>Δ</u>: ΔH/C-1 (anomeric signal) of 4,5-unsaturated 4-deoxy-L-erythro-hex-4- enepyranosyluronate; DEH: 4-deoxy-L-erythro-5-hexulosuronate hydrate; DHF: two epimers 4-deoxy-D-manno- (5*S*)-hexulofuranosidonate hydrate and 4-deoxy-D-manno-(5*R*)-hexulofuranosidonate hydrate; α/β: M/G at reducing ends of alginate residue; poly: M/G in a polymer; GM-/MG-: Alternating GM/MG polymer. Underlined labels indicate the residue with the anomeric H/C-1 giving rise to the signal, and -xx- indicates signals within a polysaccharide chain.

**Supplementary Figure 11.**
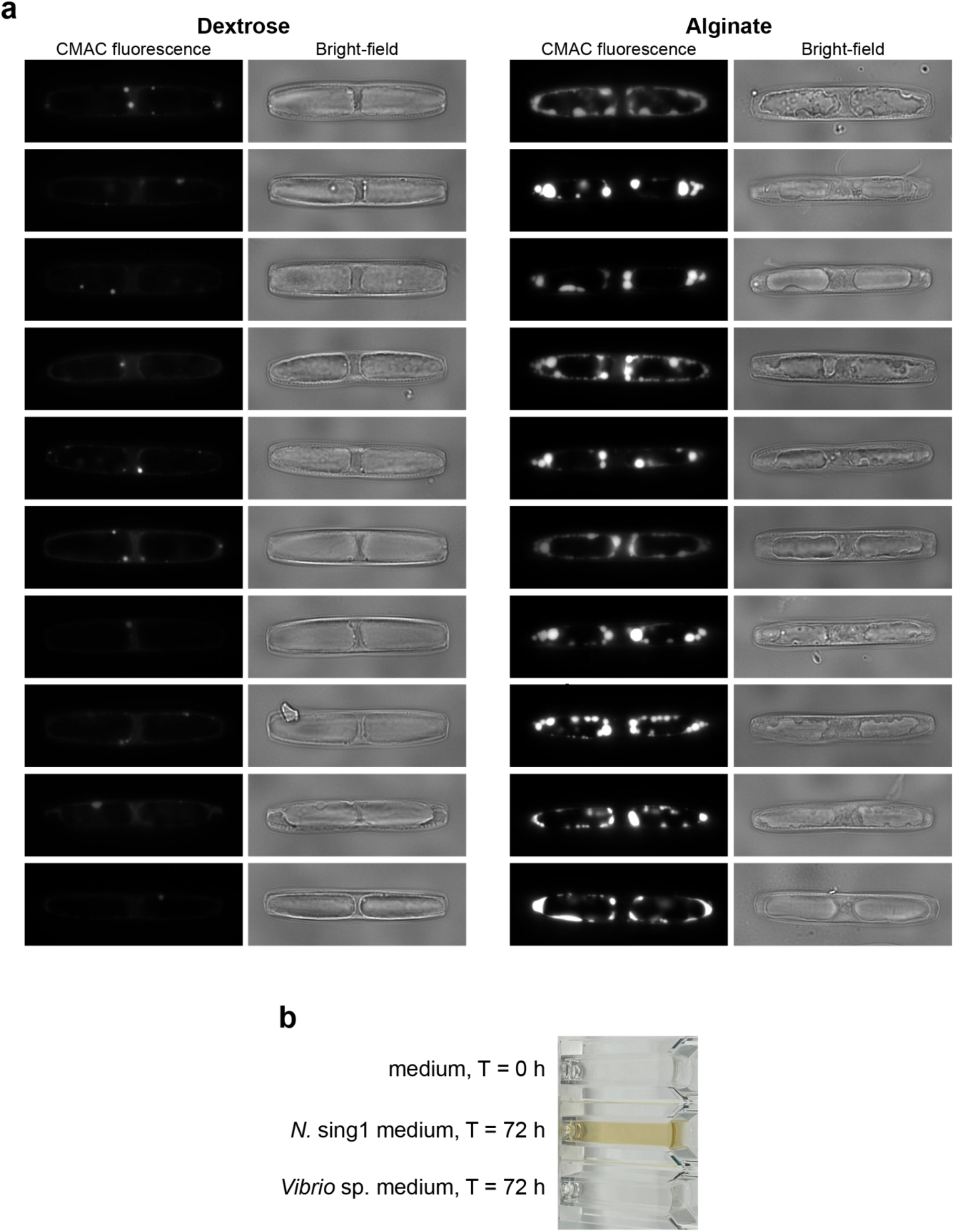
**Cell biology of diatom alginate metabolism. a**. Vacuole staining in diatoms grown on dextrose and alginate. CMAC dye shows an accumulation of vacuoles in diatoms grown on alginate, but not on dextrose (n = 10). **b**. Culture medium undergoes yellowing over time in *N*. sing1 cultures (middle, 72 h), but not in *Vibrio* sp. culture (bottom, 72 h).

**Supplementary figure 12.**
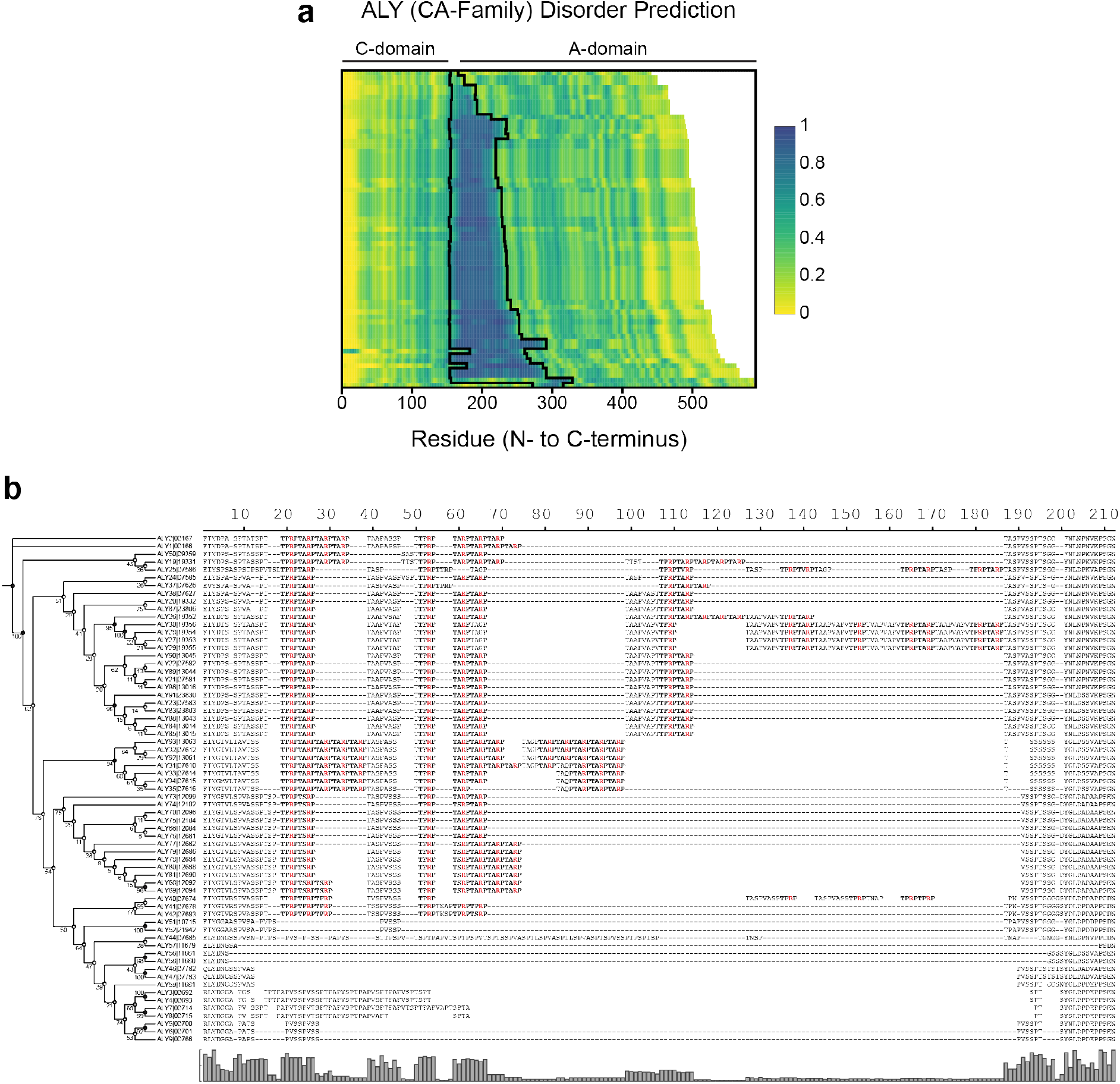
**Sequence analysis of TARP repeat sequences. a**. The heatmap shows the predicted disorder scores for *N.* sing1 ALY proteins from the CA family (using IUPred3 software). Scores are scaled as indicated (low to high disorder; green to blue). Note that the disordered regions are variable in length and occur between the C- and A-domain. The black lines denote the junction between TARP domains and folded C- and A-domains. **b**. Protein sequence alignment and maximum likelihood phylogeny of ALY (CA family) TARP repeat region sequences (bootstrap replicates, n = 1000). The alignment was manually generated to highlight how TARP repeats vary in tetrapeptide unit lengths between closely related ALYs. TARP repeats are bolded and arginine residues are highlighted in red.

**Supplementary Figure 13.**
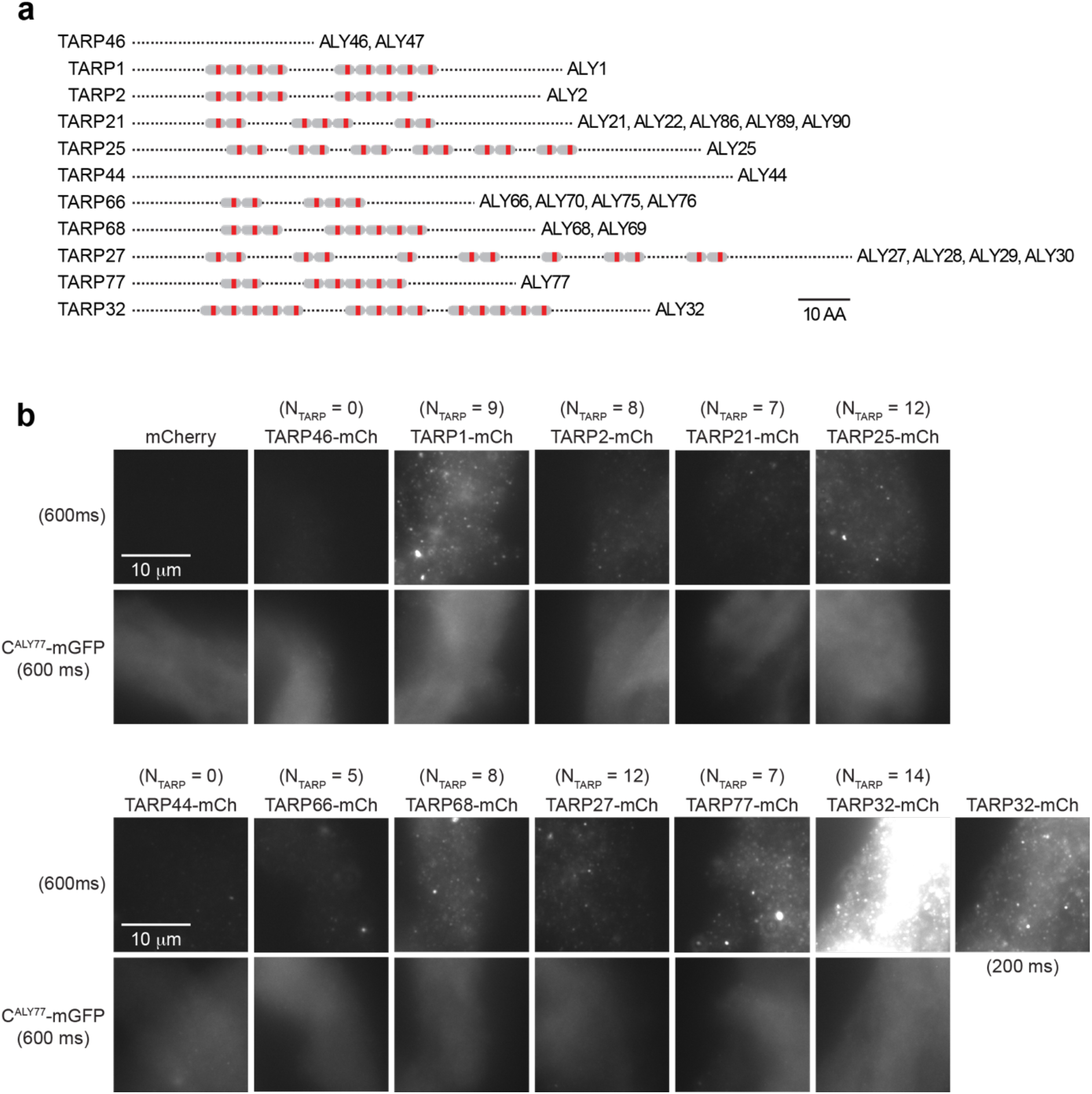
**TARP repeat sequences bind to alginate hydrogels. a**. Schematic diagram illustrating the TARP repeat units from a set of representative linkers from *N*. sing1 CA family ALY protein sequences. Each TARP repeat is annotated in gray while the red line represents the arginine residue. **b**. Double staining of alginate hydrogels with recombinant TARP domain-mCherry and C^ALY77^-mGFP fusions. NTARP denotes the number of TARP repeats in the repeat region. Brighter signals are observed in recombinant proteins that contain longer TARP repeat lengths. Samples were imaged at 100× magnification (600 ms exposure) using the TRIT-C (for mCherry; top row) and GFP (for mGFP; bottom row) channels (0 low - 800 high). For TARP32-mCh, the image was also captured at 200 ms exposure due to signal saturation when imaged at 600 ms exposure.

**Supplementary Figure 14.**
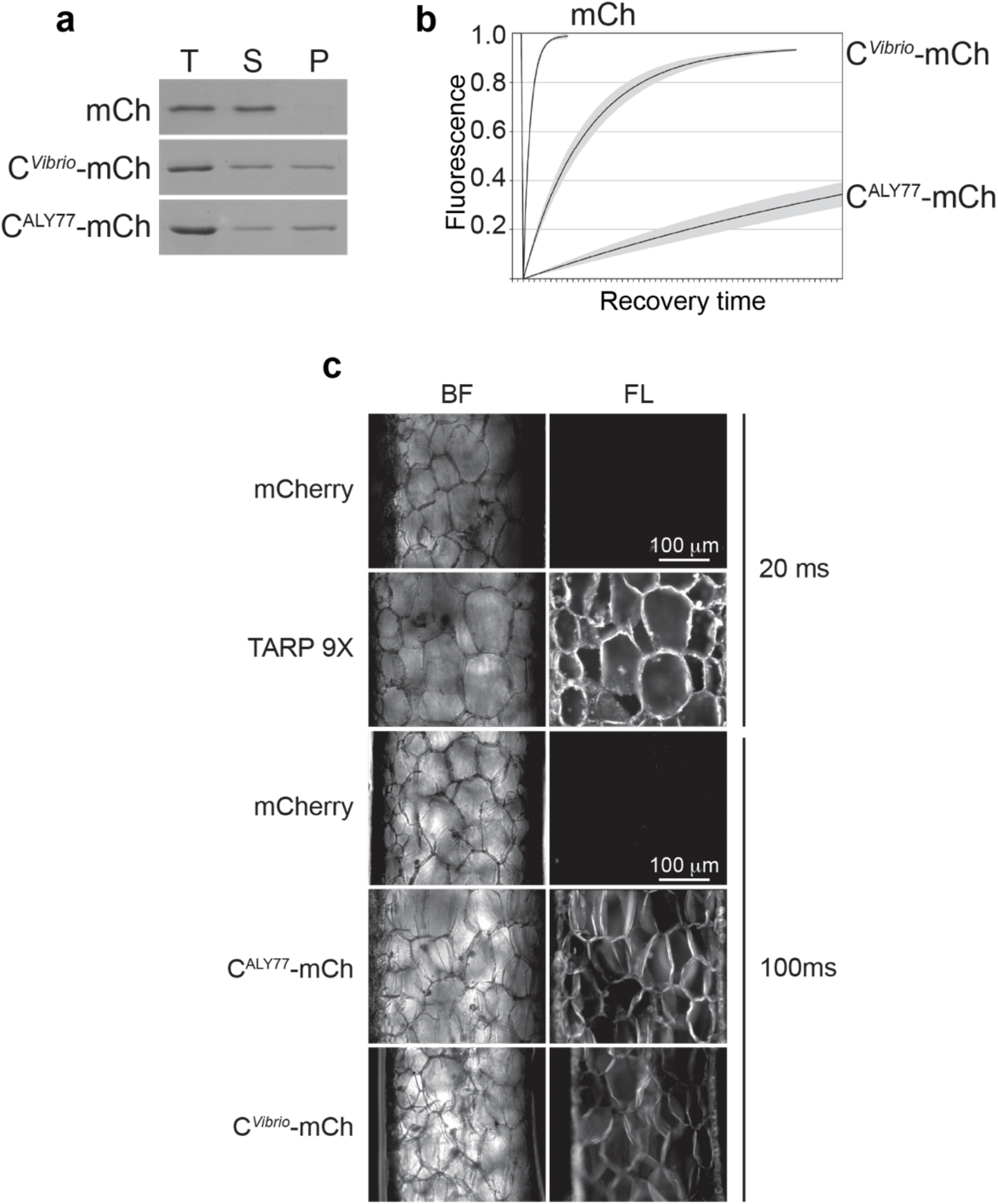
**Alginate-binding assays for CBM32 and TARP recombinant proteins. a**. SDS-PAGE of total (T), supernatant (soluble) (S) and pellet (insoluble) (P) fractions from the alginate pelleting assay with C^ALY77^ and C*^Vibrio^*-mCherry fusions. A protein band is observed in the pellet fraction for samples containing CBM32-mCherry fusions but not mCherry, indicating that the CBM32 protein binds to alginate. **b**. Fluorescence Recovery After Photobleaching (FRAP). Recovery time is slower following bleaching of the CBM32-mCherry fusion proteins as compared to the mCherry protein alone, indicating that the CBM32 proteins associate with the alginate gel. A stronger association for C^ALY77^-mCh compared to C*^Vibrio^*-mCh is observed. **c**. *Sargassum* tissue binding assay using CBM32 and TARP 9X-mCherry fusion proteins. All three fusion proteins bind to the cell wall of *Sargassum* tissue samples, which were dissected and imaged along its cross-section. Signal outlining the cell wall can be observed when tissues are visualized under epifluorescence (TRIT-C channel). No signal was observed in the mCherry control. For TARP 9X-mCh, the image was captured at 20 ms exposure due to signal saturation when using default (100 ms) settings.

## Notes

### Competing Interest Statement

The authors have declared no competing interest.

